# Multimodal X-ray imaging reveals hierarchical fibre mechanics

**DOI:** 10.1101/2025.09.19.677294

**Authors:** Elis Newham, Alissa L. Parmenter, Jishizhan Chen, Catherine Disney, Tim Snow, Joseph Brunet, Valentin Vinci, Jakub Drnec, Michael J. Sherratt, Judith A. Hoyland, Alexandre Bellier, Brian K. Bay, Nick Terrill, Peter D. Lee, Himadri S. Gupta

## Abstract

Fibrous materials—ranging from connective tissues to engineered composites—are vital to many biological and man-made systems, optimised to withstand complex in-operando or in-vivo loading. The spine’s intervertebral disc’s (IVD) load-bearing capacity depends on a hierarchical extracellular matrix, where plywood-like lamellae of collagen fibres in the annulus fibrosus contain nanometre-scale fibrils built from staggered triple-helical monomers. How intact IVDs couple fibril-scale mechanics to fibre-scale organisation under load remains unresolved. Here we introduce *TomoSAXS*, a full-field 3D small-angle X-ray scattering tomography that maps fibril-to-fibre mechanics across an intact tissue. We show that intrafibrillar molecular pre-strain (D-period stagger) is lamellar textured and tightly correlated with microscale fibre strain. Pre-strain is inversely related to fibril strain and its variability, consistent with load-sharing through molecular unwinding. Radial strain bridges and high-curvature zones at the annulus fibrosus–nucleus pulposus interfaces emerge as critical regulators of local mechanics. These findings reveal concerted fibril–fibre interactions that sustain mechanical equilibrium in the IVD, preserving elasticity and shape. More broadly, *TomoSAXS* establishes a platform to visualise nano- to micro-scale matrix mechanics across biological and synthetic fibrous materials, with applications in ageing and disease, therapeutic evaluation, and the design of bio-based and bioinspired materials.

## Introduction

Fibrous materials provide critical functions across biological and engineered systems. Biological tissues achieve remarkable properties through hierarchical structuring from the nano-/molecular to whole-organ-scales, offering design principles for advanced synthetic materials^1,2^, such as nanostructured polymer composites^3^. Collagen-rich tissues such as tendon, cartilage, and bone underpin vertebrate biomechanics, with function emerging from complex, hierarchically structured assemblies of collagen fibrils (typically Type I or II)^4–6^ and non-collagenous components, including hydrated glycosaminoglycans^7^. Distinct features at each hierarchical level, such as nanoscale pre-strain gradients^8^ and microscale variations in fibre orientation and curvature^9^, contribute to tissue performance. However, the complex architecture across scales presents major challenges in isolating the respective effects of each hierarchical element and understanding how they collectively support tissue function.

A clinically important example is the intervertebral disc (IVD), a sophisticated composite structure in the spine which must balance flexibility with efficient distribution of load^10^. IVD injury and age-related degeneration disrupts the mechanical environment and is a key driver of back pain – the leading global cause of years lived with disability and a significant economic burden^11^. While emerging therapies such as synthetic disc replacements hold promise^12^, their development is hindered by an incomplete understanding of IVD structure– function relationships across length scales.

Here we introduce *TomoSAXS*, a correlative, multiscale X-ray imaging approach that integrates phase-contrast computed tomography (pCT)^13^ at the microscale with small-angle X-ray scattering (SAXS)^14^ at the nanoscale. We apply this technique to intact IVDs under physiologically relevant loading to investigate structure–function relationships across multiple length scales.

The IVD consists of a central nucleus pulposus (NP) – a hydrated proteoglycan rich matrix that redistributes mechanical loads – surrounded by the annulus fibrosus (AF), a fibrous outer ring which provides structural support and stability^15^ (Figure 1ai). Within the AF, concentric lamellae tens of microns thick are composed of collagen fibres (1-20 μm diameter) with alternating orientations between adjacent lamellae (Figure 1aii), forming an architecture believed to convert NP-derived hydrostatic pressure into circumferential tensile stress^10,15^. An inter-lamellar network, composed of a non-fibrillar matrix, elastic fibres, and trans-lamellar cross-bridges, further stabilises this structure by anchoring adjacent lamellae^16^.

Imaging of this fibrous microstructure across the whole organ is challenging due to the similar density of fibres and the surrounding matrix. However, the phase information revealed by pCT, combined with digital volume correlation (DVC), is uniquely suitable to resolve the low-contrast fibre structure (Figure 1c) and volumetric in-situ deformation and strain at the microscale. Recent applications of DVC have revealed intricate, region-specific strain patterns, with tissue compliance linked to fibre curvature and orientation^9^. These complex patterns depend crucially on the intrinsic structural properties of the fibres.

**Figure 1.**
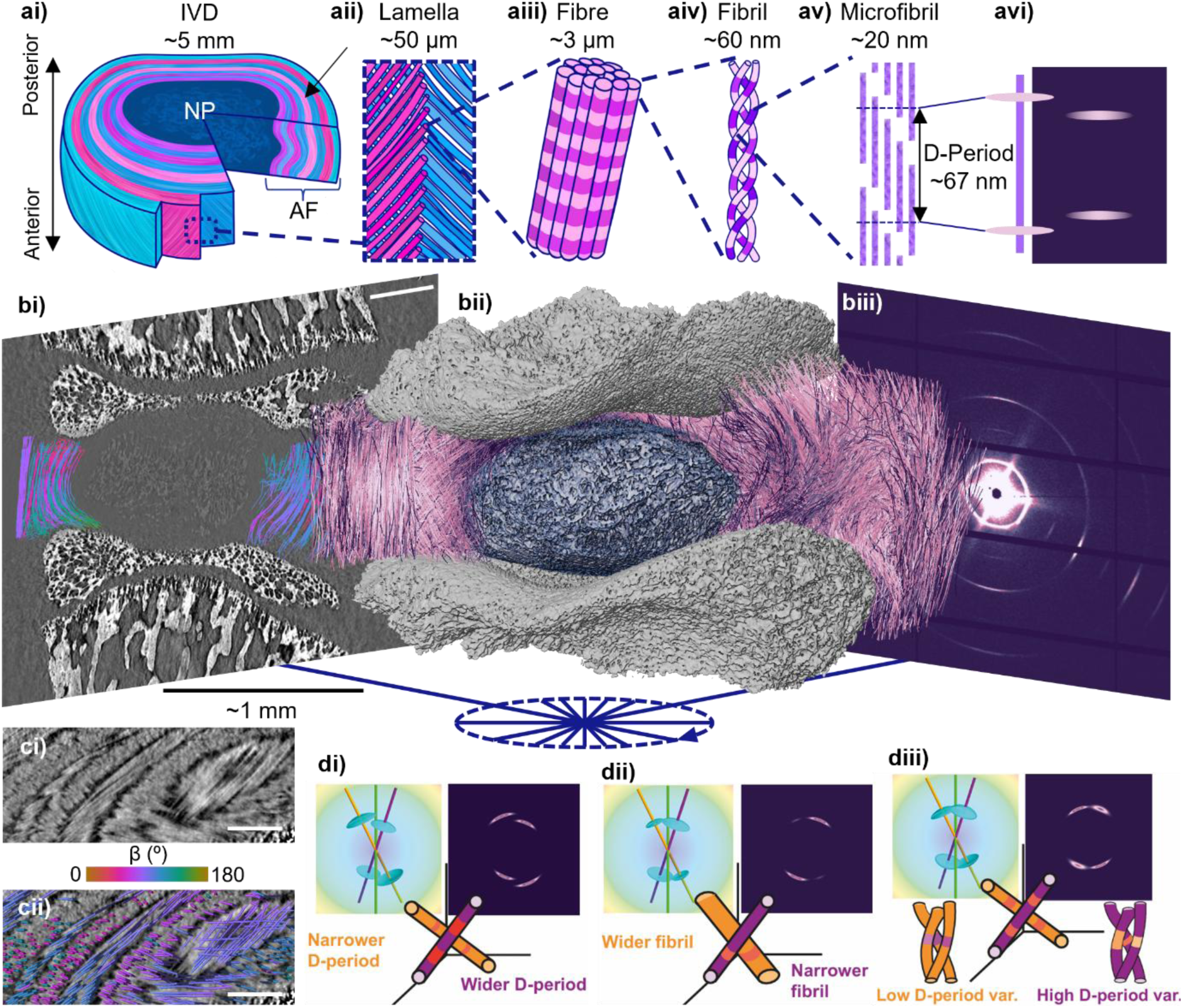
Hierarchical structure of the IVD and correlative pCT and SAXS measurements. a) Illustration of the hierarchical structure of collagen in the IVD; ai) macroscale structure of the annulus fibrosus (AF) and nucleus pulposus (NP), aii) lamellar plywood-like structure of collagen fibres in the AF, aiii) a collagen fibre consisting of multiple fibrils, aiv) a collagen fibril formed of helically wound microfibrils, av) a microfibril of staggered tropocollagen molecules with D-period labelled, avi) schematic of X-ray scatter produced by collagen D-period. b) Multimodal X-ray imaging of the IVD; bi) slice through the pCT image with traced fibres overlayed and coloured by lateral angle β (scale-bar represents 0.6 mm), bii) 3D rendering of the IVD as captured by pCT, with AF fibres rendered pink, NP rendered blue, and vertebral endplates rendered in grey, biii) resultant SAXS intensity pattern from an X-ray beam-path through the IVD. c) Fibre tracing of individual collagen fibres, ci) greyscale pCT slice showing clear fibrous microstructure, cii) traced fibres overlayed on the pCT image coloured by β angle (scale-bars represent 300 µm). d) Schematic examples of nanoscale fibril properties reflected in SAXS patterns, di) D-period, related to peak position along the radial axis, dii) fibril diameter, inversely related to peak width along the circumferential axis, diii) D-period variation, inversely related to peak width along the radial axis.

At the nanoscale, collagen fibres are composed of fibrils (∼20-100 nm, Figure 1aiv) which are in turn formed of microfibrils of staggered tropocollagen molecules with a mean periodicity (D-period) of ∼65-67 nm ^4^ (Figure 1av). This axial staggering gives rise to characteristic SAXS diffraction patterns with Bragg peaks at integer multiples of 1/D (Fig 1avi), which act as an internal gauge of fibrillar strain, orientation, and diameter (Figure 1d). In other fibrous tissues – such as bone, tendon, and cartilage – two-dimensional (2D) transmission-based SAXS has been used to identify nanoscale deformation mechanisms, including fibrillar pre-strain gradients and hydration-induced alteration in diameter in cartilage^8^, nanoscale response to *in situ* loading in tendons^17^, and altered mineral-collagen shear-transfer in ageing bone^18^.

Linking 3D deformation of fibrous structures across scales using SAXS and pCT poses major challenges. Standard SAXS analysis is based on 2D convolutions of combined 3D reciprocal scattering from multiple fibres encountered along individual X-ray beam-paths. Since fibre scattering is anisotropic, the 2D pattern depends on fibre orientation at each scattering voxel^8^, invalidating traditional scalar tomographic methods for macroscopic specimens with complex fibre texture^19^. To overcome this, tensor tomographic methods have been developed which use multi-axis rotation and spherical basis functions to reconstruct 3D nanostructure in a voxel-wise manner in undeformed structures^20,21^. However, there remains an unmet need for methods to reconstruct both micro- and nanostructure of individual fibres and track their heterogenous deformation within the broader context of micro- to macroscale analysis.

Here, *TomoSAXS* uses fibre-tracing of collagen fibres in the AF from pCT reconstructions and their volumetric registration with tomographic SAXS data to allow deconvolution of the 3D scattering from individual fibres in each SAXS beam-path. The IVD is an excellent biological exemplar, due to its lamellar fibrous texture allowing deconvolution of contribution from different fibrils under rotation. Instances through the SAXS tomography where fibres are predicted to provide sufficient portions of independent scattering intensity are sampled and fitted with 3D fibre-diffraction models to estimate their individual nanoscale properties (mean and variation in D-period, fibril diameter). Registration between SAXS and pCT volumes allow these to be volumetrically mapped onto pCT reconstructions, alongside DVC measurements of tissue and fibre deformation. The main advantages of this technique are: a) the isolation of nanoscale properties of individual collagen fibres; b) the estimation of per-fibre nanoscale strain through tracking the location of fibres in their unloaded and loaded states; and c) the spatial mapping of nanoscale properties facilitating hierarchical structural and mechanical analysis from the nano- to macroscales. This reveals new insights into hierarchical IVD mechanics, providing an important case study for the coupled nano-microscale characterization of other collagenous tissues, such as arteries and cartilage, and engineered fibrous materials, such as carbon fibre composites. *TomoSAXS* opens new avenues for linking collagen structure to function in health, disease, ageing and tissue engineering, and for advanced characterisation of next-generation composites from biomedical to aerospace applications.

**Extended Data Figure 1.**
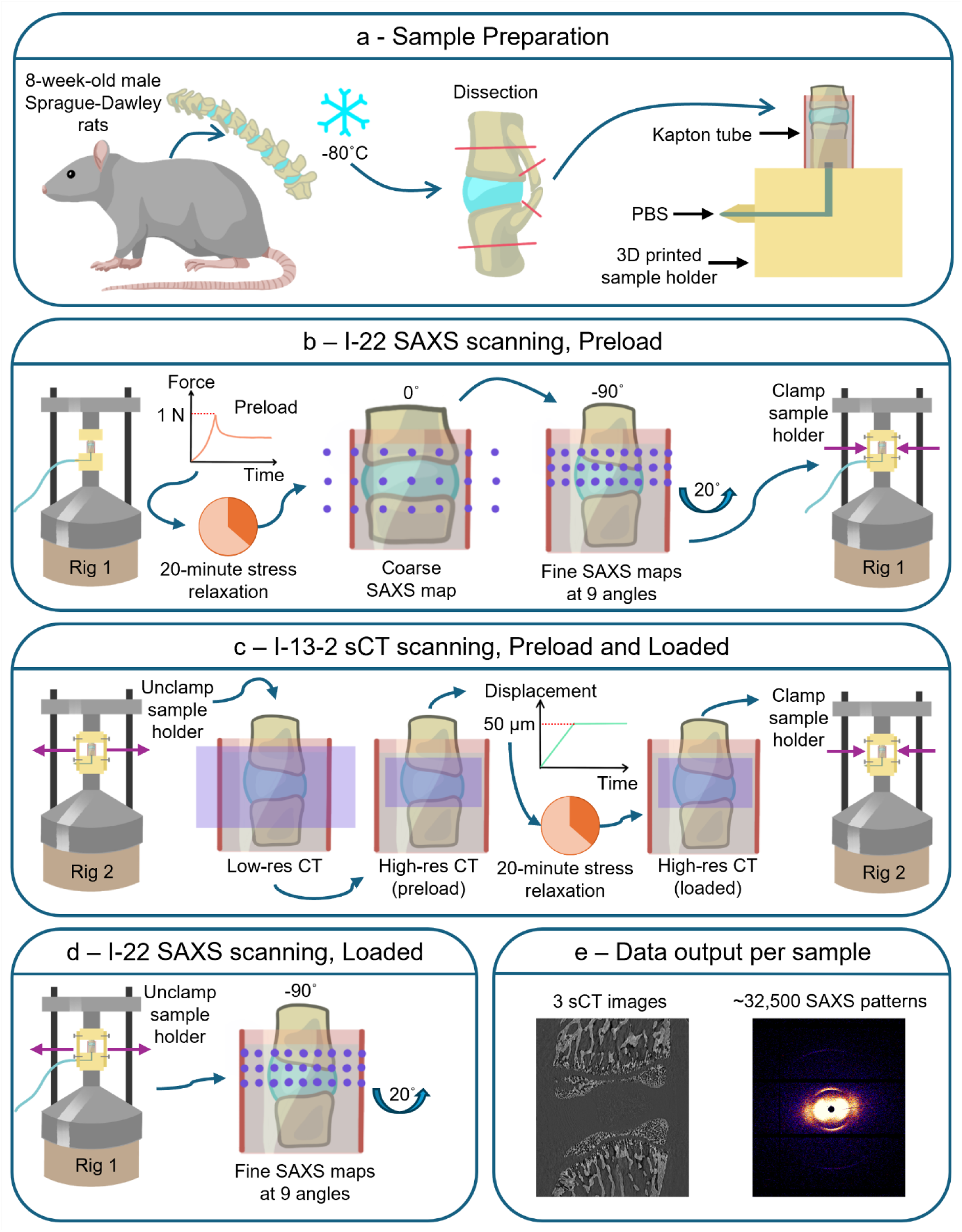
*in situ TomoSAXS* experimental method. Illustrations of a) sample preparation and sample environment, b) preload application and SAXS scanning, c) pCT scanning before and after compressive displacement, d) post-loaded SAXS scanning, and e) summary of experimental data output per sample.

### *TomoSAXS*: Experimental Methodology and 3D SAXS reconstruction

To obtain multiscale biomechanical analysis of IVDs under physiologically relevant load-levels, correlative pCT and single axis tomographic SAXS were performed at the Diamond Light Source synchrotron on intact murine IVD samples before and after the application of 50 μm compressive displacement (Methods). An illustration of the experimental method is shown in Extended Data Fig. 1. *TomoSAXS* requires the indexing of fibres and characterisation of their orientations from a 3D image, achieved here through automated fibre-tracing of the initial pCT image^9^ (Fig. 1c) (Methods). This orientation data, alongside rotational SAXS sampling and spatial registration of pCT and SAXS datasets (Methods), is key to deconvoluting the individual contributions of fibres across SAXS X-ray beam-paths, as SAXS signals are orientation dependent. Unlike SAXS tensor tomography^19^, where multiple tomographic axes are required for a full reconstruction, *TomoSAXS* uses a single axis for SAXS tomography. The isolation of individual fibres from pCT enables the reconstruction of SAXS parameters on a per-fibre basis, allowing spatial mapping of nanoscale properties at the resolution of the pCT images (here 1.6 µm voxel size, 5.5 µm resolution^22^) as opposed to being limited to the SAXS voxel size (here 20 µm). This is vital for complex fibrous materials like the AF, where rapid angular changes occur at the microscale. Fibre tracing also minimises the need for repeated angular sampling during SAXS scanning (resulting in significant X-ray doses to biological samples^23^), and a single axis of rotation is applied to maximise the angular independence of scattering from different fibres rather than to estimate their orientation.

A typical SAXS beam path contains multiple collagen fibres with varying orientation (Fig. 2a); however, due to their 3D orientation and anisotropic diffraction, only a subset of these fibres will satisfy the Ewald diffraction condition and visibly contribute to the SAXS pattern (Fig. 2b-c). The first step of the *TomoSAXS* reconstruction uses Singular Value Decomposition (SVD) of 3D fibril diffraction modelling to estimate the scattering amplitude of individual fibres in each beam path (Fig. 2c), using predicted mean values for physical properties to provide an initial amplitude estimation (Methods).

**Figure 2.**
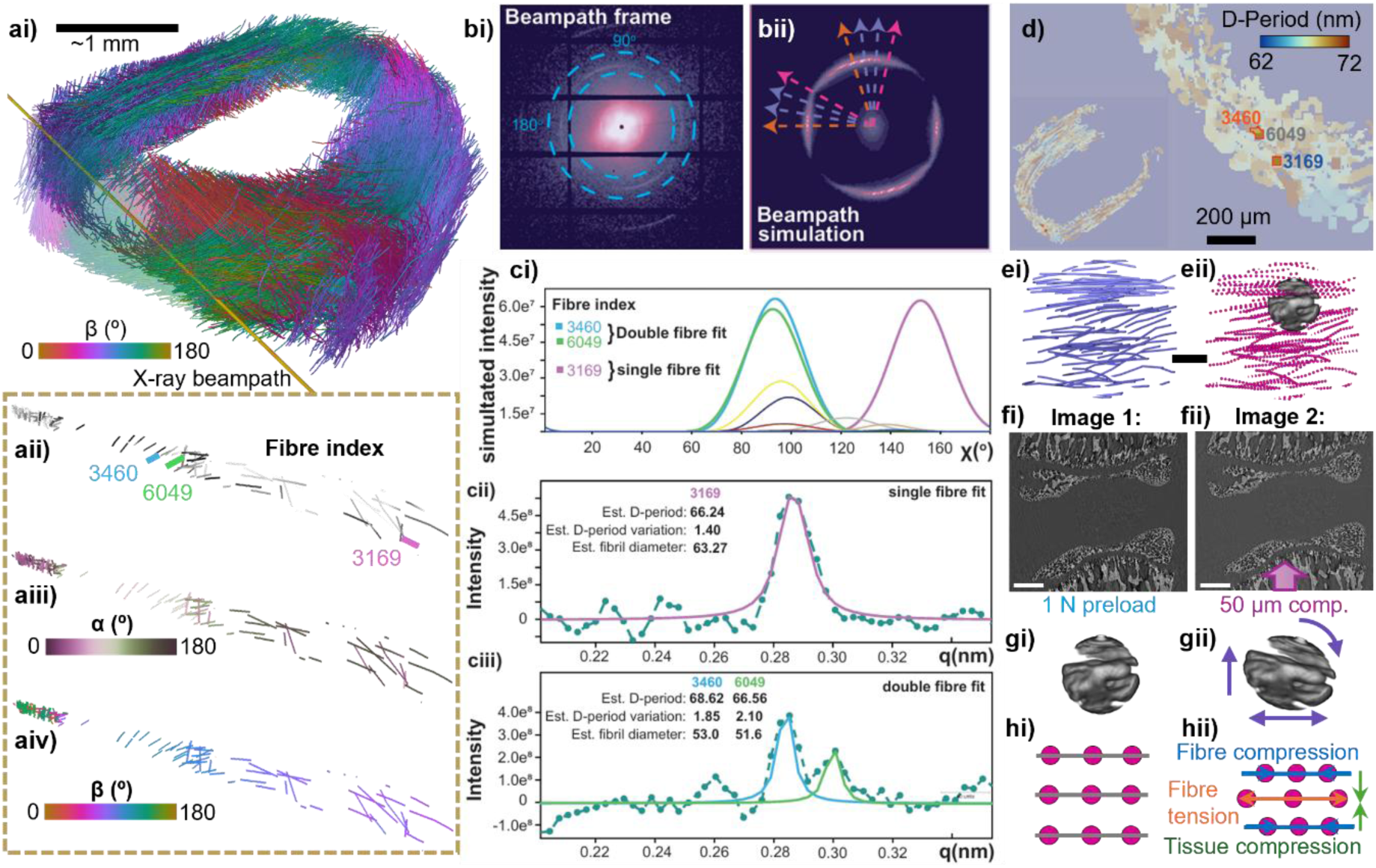
*TomoSAXS* reconstruction and fibre-based DVC. ai) Complete fibre tracing for an IVD sample, with fibres coloured by orientation and one SAXS beam path rendered in gold, aii) fibre index for the three fibres predominantly contributing to SAXS signal along the beam path, aiii) α and aiv) β orientation angles for the fibres withing the beam path. b) Resultant bi) real and bii) simulated SAXS frames for the beam path represented in (a). c) Fitting and estimation of nanoscale fibril properties from isolated scattering regions. d) Spatial reconstruction of per-fibre nanoscale properties in the pCT cross-section; the locations of the three fibres from a)-d) indicated. e) Point cloud generation from fibre tracing, with the size of one sub-volume rendered in grey, scalebar represents ∼65 µm; f) pCT images after i) 1 N preload and ii) 50 µm compressive displacement, scalebars represent 300 µm; gi) 3D rendering of a sub-volume used for DVC tracking between images, gii) sub-volume after translation, rotation, and first order strain to match the deformed image; h) illustration of point cloud i) before and ii) after compression, demonstrating different modes of tissue and fibre deformation.

Next, the 3D orientation and estimated scattering amplitude of fibres are used to perform an iterative simulation of the SAXS tomography and isolate angularly independent scattering instances (Methods). These are isolated when one, two, or a maximum of three fibres contribute predominantly (>90%) to total scattering across at least 10° around the circumferential χ axis of individual SAXS frames (Fig. 2ci). By sampling SAXS intensity within these regions across three sub-sections of equal angular width along the *q* (radial) axis and fitting profiles with 3D diffraction models^8^, the physical parameters fibril D-period, D-period variation, and fibril diameter (Fig. 1d) can be obtained for each fibre (Fig. 2cii-iii) (Methods). Reconstruction follows a cascading pattern, where the solution of individual fibres enables further deconvolution of neighbouring fibres, allowing an iterative sweep through the sample until every possible fibre has been fitted. Fibril properties can then be mapped onto the pCT image, enabling assessment of spatial variation and gradients in nanoscale properties (Fig. 2d, Methods).

The measurement of strain in per-fibre nanoscale properties requires accurate tracking of fibre deformation during *in situ* experimentation; achieved here with DVC analysis^9^. DVC analysis was performed on pCT datasets to measure the deformation of collagen fibres in the AF following axial compression using a local, sub-volume based approach, with measurement points placed along the traced fibres (Fig. 2e-f, Methods). Application of DVC-measured displacements to traced fibres in their un-deformed state allows calculation of their location and orientation following loading, while preserving fibre index. This information is used to reconstruct SAXS data of loaded fibres and compare resultant fibrillar parameters across loading to generate nanoscale strain values. Fibre and tissue level strains can also be obtained from the DVC-measured displacements. Fibre strain is calculated from a weighted polynomial fit to the displacements along the fibre direction (Fig. 2f, Methods). At the tissue level, the Lagrangian strain tensor for each point is calculated from a polynomial fit to the displacement field.

**Extended Data Figure 2.**
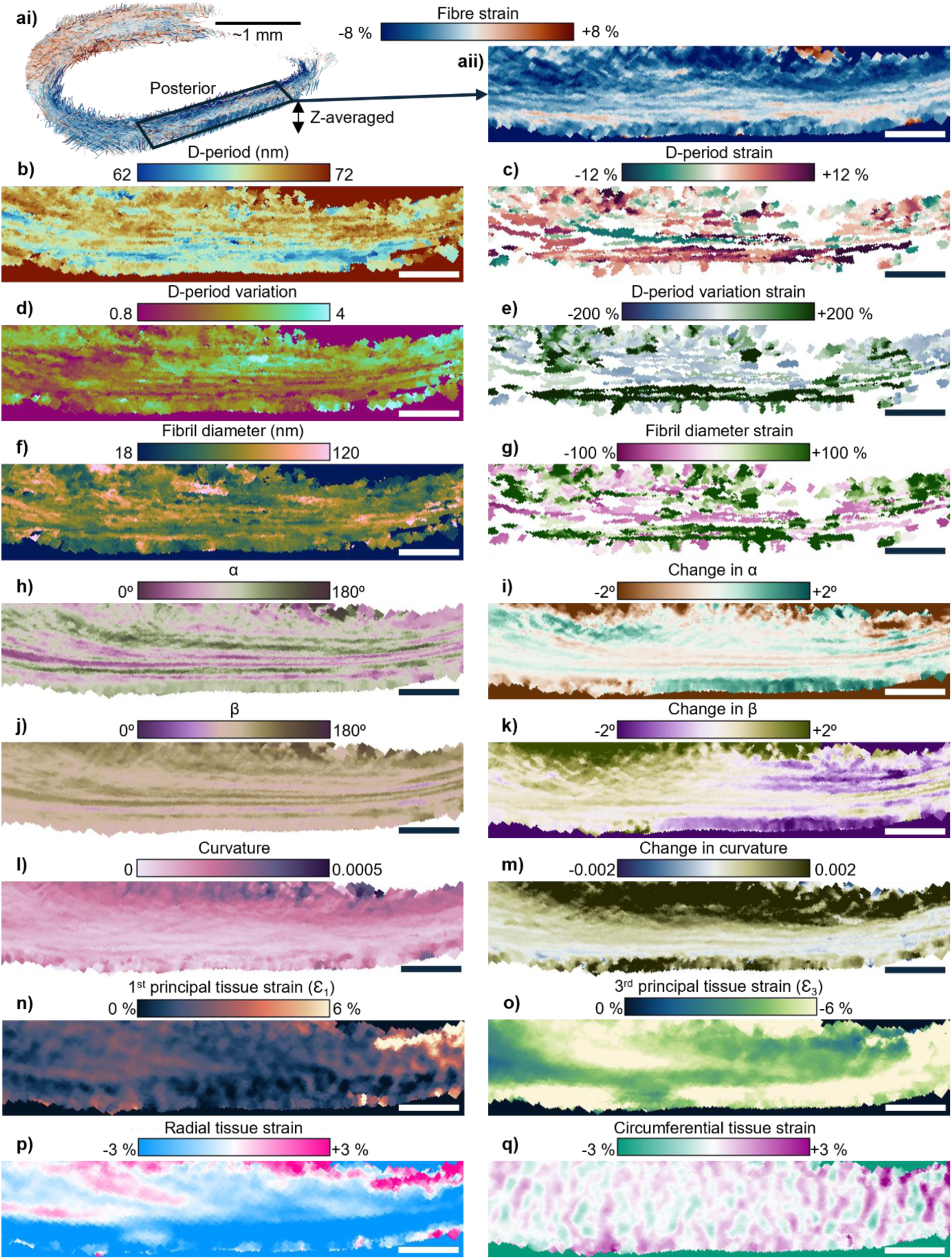
*TomoSAXS* enables correlative spatial mapping of nano- to microscale structure and mechanics. ai) Fibre strain in the AF, aii) z-averaged fibre strain in the posterior region as highlighted in ai. Z-averaged spatial mapping of: b) per fibre mean D-period, c) per fibre mean D-period strain – the percentage change in D-period measured before and after loading, d) per fibre D-period variation, e) per fibre D-period variation strain, f) fibril diameter, g) fibril diameter strain, h) vertical angle α, i) change in α with load, j) lateral angle β, k) change in β with load, l) fibre curvature, m) change in curvature with load, n) 1^st^ principal (maximum tensile) tissue strain, o) 3^rd^ principal (maximum compressive) tissue strain, p) radial tissue strain, q) circumferential tissue strain.

### *TomoSAXS* enables nano- to macroscale analysis of fibrous materials

*TomoSAXS* correlates 3D microscale imaging with nanoscale measurements from SAXS, enabling 3D mapping and direct statistical comparison of per-fibre micro- and nanoscale properties in macroscopic samples. Here, application of *TomoSAXS* has resulted in 3D maps of structural and mechanical properties specific to individual collagen fibres across the spatio-structural hierarchy of the AF (Fig. 3; Extended Data Figs 2-3). The orientational texture of our samples and applied experimental parameters (Methods), permitted nanoscale reconstruction for >55% of all fibres. This provided sufficient resolution to highlight complex textural patterns in both structural and mechanical nanoscale properties (Fig. 3c,e, Extended Data Fig. 2b-g).

**Figure 3.**
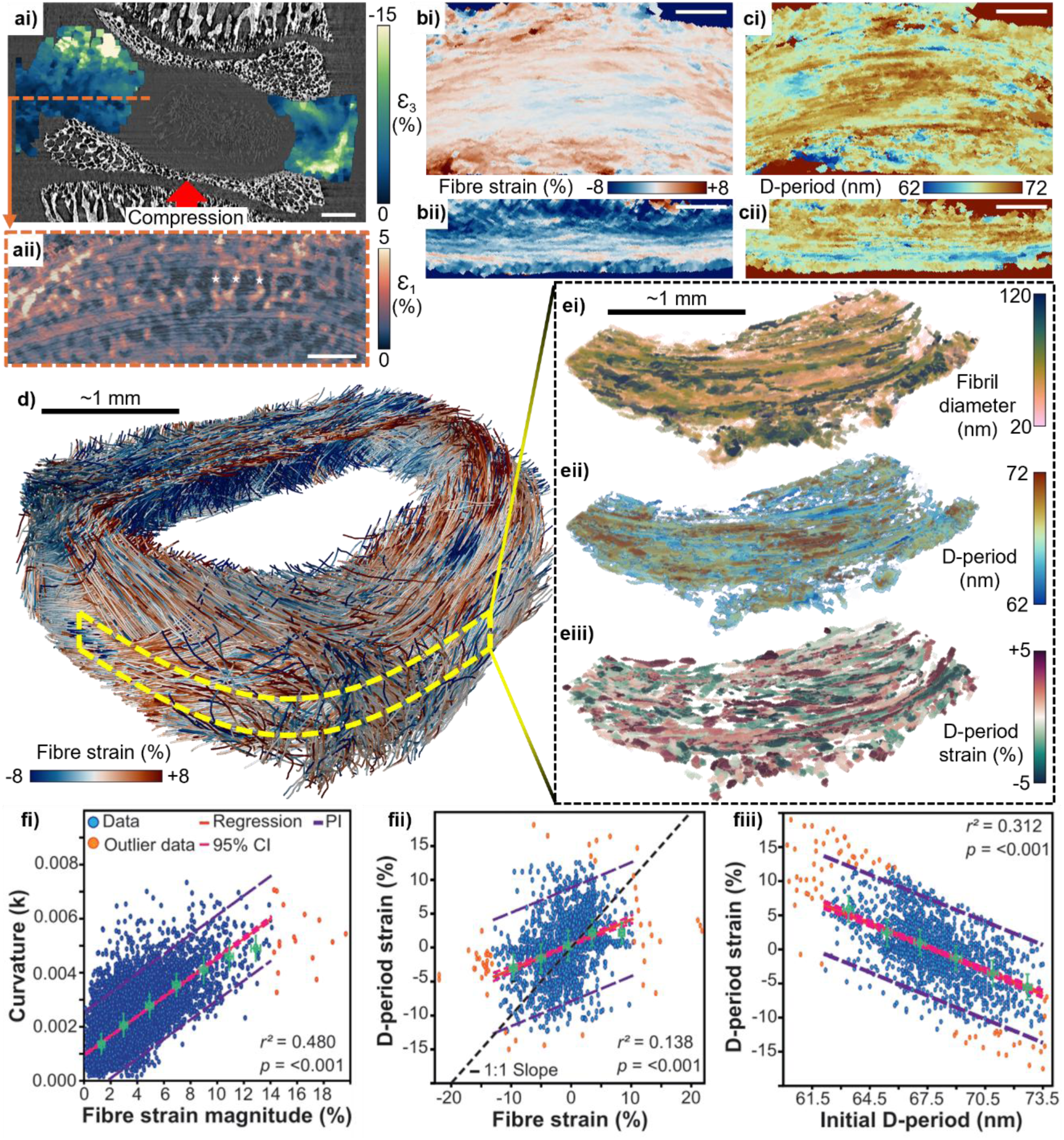
*TomoSAXS* reveals multiscale collagen structure–function relationships in whole intervertebral discs. a) Tissue-level patterns in ai) compression (3rd principal strain ε_3_) and aii) tension (1st principal strain ε_1_). b) Microscale fibre strain patterns in the bi) anterior and bii) posterior annulus fibrosis (AF). c) Nanoscale patterns in initial D-period showing anti-correlation with fibre strain patterns in (b). d) 3D reconstruction of fibre strain patterns through the entire AF. e) Maps of nanoscale properties found within the region in (d) marked with yellow dashed lines. f) Regression analysis of fi) fibre curvature vs fibre strain magnitude, fii) fibrillar D-period strain vs fibre strain, fiii) fibrillar D-period strain vs initial D-period. 95% CI (pink dashed lines): 95% confidence interval; PI (purple dashed lines): prediction interval.

An estimated 100 nm accuracy in DVC fibre-displacement tracking (Methods) allowed confident estimations of displacement under load for individual fibres, enabling precise per-fibre nanoscale D-period and fibril diameter strain measurements (Fig. 3eiii, Extended Data Fig. 2c,e,g). The nanoscale measurements of structure and mechanical response can then be directly compared with those at the fibre and tissue levels (Fig. 3, Extended Data Fig. 2).

### Tissue-level strain patterns reveal a mechanical role for non-collagenous matrix in the AF

At the highest level (Fig. 3ai-ii), local tissue strains measured from DVC were overall higher than the bulk strain applied to the IVD, measured as the percentage change in disc height (Methods, Supplementary Note 8). A reduction in IVD height of 2.2 % resulted in tissue level principal compressive strains of 4.2 ± 3.3 % in the AF, with tissue level strains greater than 15 % observed particularly in regions close to the vertebral endplates (Fig. 3ai). These high tissue level strains agree with previously reported values in human IVD under compressive load^24^, and local tissue strains higher than the overall applied strain have been observed in other hard-soft tissue junctions^25^.

Maps of tissue-level 1^st^ principal (tensile) strain reveal both lamellar and trans-lamellar patterns (Fig. 3aii). Projecting tissue level strain along the circumferential direction (Extended Data Fig. 3a) shows alternating trans-lamellar bands of tension and compression (Extended Data Fig. 3c). We hypothesise these strain patterns may be related to the trans-lamellar cross-bridging network of the AF (Extended Data Fig. 3b), as previously suggested^26^, which is thought to aid in returning AF tissue to its original shape after deformation^16^. Because strains were recorded after stress relaxation, these trans-lamellar patterns may be a result of elastic recoil from the inter-lamellar matrix. Such patterns suggest that trans-lamellar cross-bridges may be an initiation site for annular tears, providing support for previous hypotheses^27^. Spatial mapping of strain in the radial direction shows significant radial compression of the AF due to bulging of the NP (Extended Data Fig. 3d), with some lamellae exhibiting radial tension, possibly due to swelling induced by fluid flow out of the NP into the AF.

Tissue-level principal strain direction, measured as the angle between the strain eigenvector and the local fibre tangent (Extended Data Fig. 3g), averaged 55° for compression and 58° for tension (Extended Data Fig. 3e), indicating substantial strain accommodation by the extrafibrillar matrix and inter-fibre motion. Spatial mapping of principal strain directions shows clear lamellar texturing, with alternating lamellae demonstrating tissue level principal strains acting parallel or orthogonal to the fibre direction (Extended Data Fig. 3f,h).

**Extended Data Figure 3.**
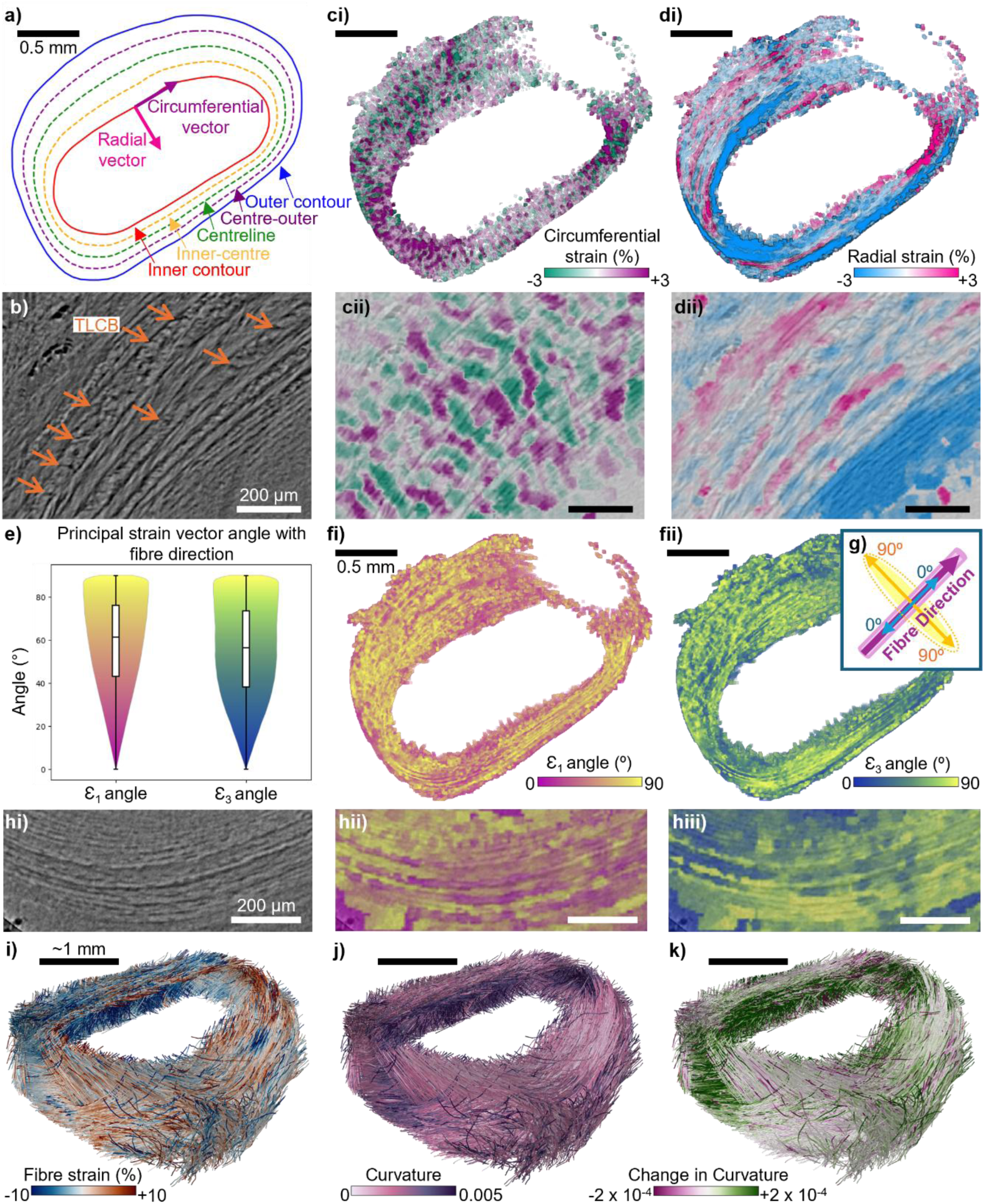
Tissue level strain analysis and the link between fibre strain and curvature. a) Contour lines used for calculating tissue-level radial and circumferential strains (Supplementary Note 6); b) greyscale image of lamellae in the anterior AF, with trans- lamellar cross-bridges (TLCBs) highlighted by orange arrows; c) circumferential and d) radial strain in the AF. c) Violin plots of 1^st^ and 3^rd^ tissue level principal strain angles to the local fibre direction; f) 3D renderings of i) 1^st^ and ii) 3^rd^ principal strain angle with the local fibre direction; g) illustration of the angles used for defining principal strain vector angle with the local fibre direction; hi) greyscale image of the posterior-lateral AF with ii) 1^st^ principal strain angle and iii) 3^rd^ principal strain angle displayed as a colourwash, showing clear lamellar texturing. 3D renderings of fibres in the AF coloured by i) fibre strain, j) curvature, and k) change in curvature upon compressive loading.

### Highly curved fibres experience the highest strains

Fibre level strain magnitude showed a significant positive correlation with fibre curvature (*r*^2^ = 0.480, *p*<0.001) (Fig. 3fi, Extended Data Fig. 3i,j). Fibres with the greatest curvature were typically located at the transition zone between the NP and AF in the posterior and lateral regions of the IVD (Extended Data Fig. 3j), with these fibres showing a tendency to increase in curvature under compressive loading of the IVD (Extended Data Fig. 3k). The link between fibre curvature and fibre strain may partially explain why the posterior and posterior-lateral regions of the IVD are most prone to herniation^28^.

### Fibrillar pre-strain drives fibre-level mechanical behaviour

Fibre strain measured from DVC showed a significant positive correlation with fibrillar D-period strain measured from SAXS (*r^2^* = 0.138, *p*<0.001) (Fig. 3fii). A mean fibre strain magnitude equal to 3.70% (±3.19%) and mean fibrillar strain magnitude equal to 3.60% (±3.36%) (Fig 4d) suggests that most of the strain at the fibre level is passed down to the fibril, with limited inter-fibril sliding at this level of loading. However, the slope of the regression line deviates from one due to a significant proportion of fibres with near-zero D-period strain which experience a considerably higher fibre strain magnitude (Fig. 3fii). This may be evidence of previously reported lags in fibril strain response to low strain conditions due to molecular tilting in twisted collagen fibres^29^.

**Figure 4.**
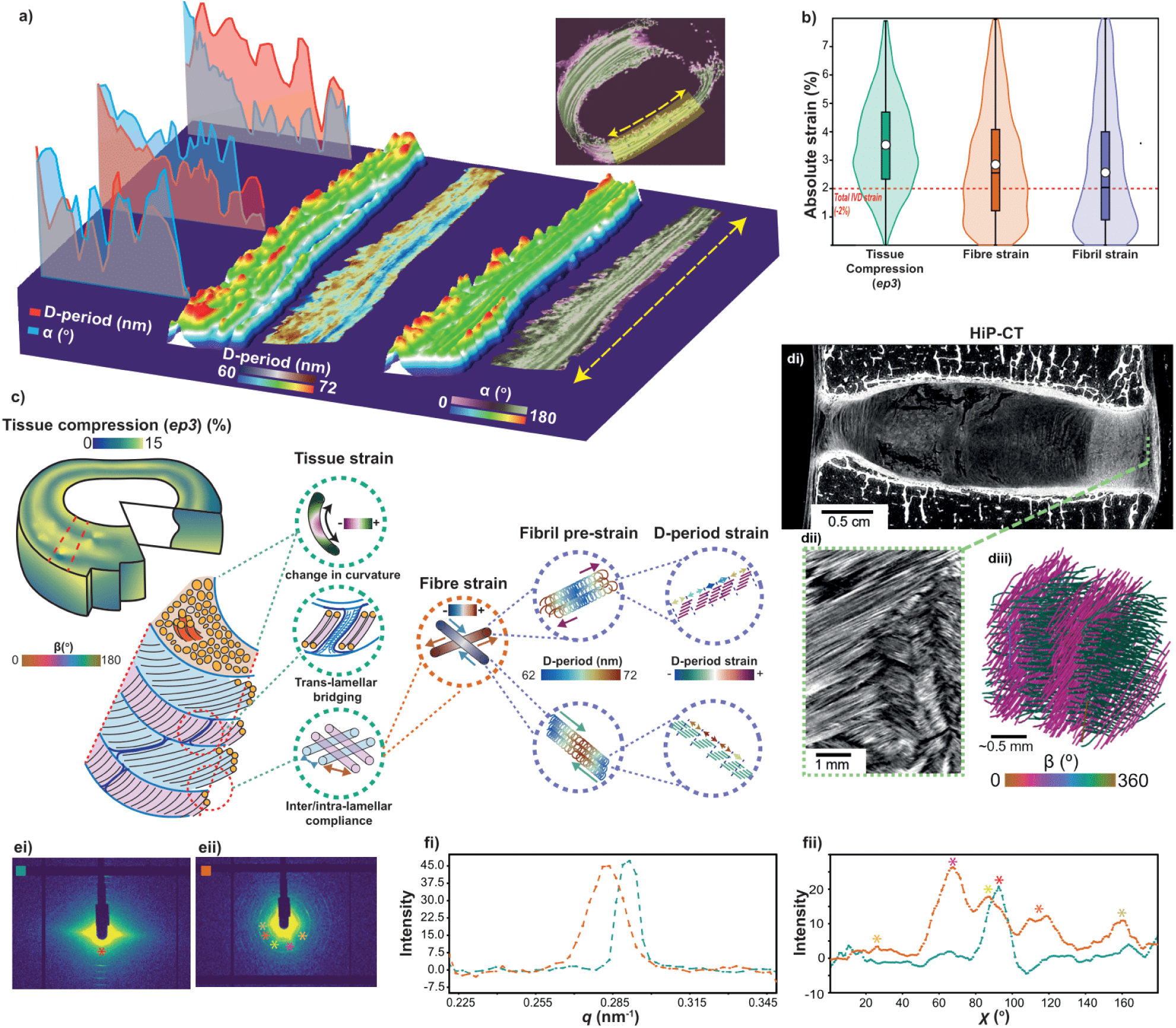
Spatial analysis of *TomoSAXS* data, hierarchical mechanics of the IVD, and *TomoSAXS* of human IVD. a) Transect analysis and comparison between D-period patterning and vertical fibre orientation (α) across algorithmically straightened samples of the posterior IVD. b) Comparison of absolute strain magnitudes between hierarchical scales in the AF. C) Schematic outlining the mechanisms of load transfer isolated here from the macro/whole tissue level, to the nanoscale. d) Preliminary *TomoSAXS* data of human IVD using HiP-CT (di,ii) and fibre tracing (diii) (hierarchical phase-contrast tomography). e) Example tomographic SAXS frames from differing rotations and locations. f) SAXS data integration across *q* (fi) and *χ* (fii).

Fibrillar D-period strain showed a significant negative correlation with initial fibrillar D-period (*r^2^* = 0.312, *p*<0.001) (Fig. 3fiii). This can be interpreted as fibres with an initially tensile pre-strain undergoing compression upon loading, and those with an initial compressive pre-strain undergoing tension. Initial fibrillar D-period covered a broad range of values, from 60 nm to 74 nm. Assuming a zero-strain state of ∼67 nm, this represents residual fibrillar strains ranging from -10 % to +10 % in the IVD. Given a Young’s Modulus of ∼1-2 GPa for hydrated Type I collagen fibrils^30^, our data translates to a measured pre-stress of 100-200 MPa. The regression line for initial D-period versus D-period strain shows that D-period strain values typically return the fibril D-period towards its zero-strain state of 67 nm.

Mean fibrillar D-period demonstrated a significant positive correlation with intra-fibrillar variation in D-period (*r^2^* = 0.089, *p*<0.001) (Extended Data Fig. 4a,b). Fibrils with larger mean D-periods, which can be considered as having a tensile pre-strain, have greater variability in the D-periods of their constituent microfibrils. As for D-period pre-strain and strain, we also find a significant negative relationship between initial per-fibre D-period variation and change in variation upon load (*r^2^ =* 0.340, *p*<0.001) (Extended Data Fig. 4), suggesting that fibres undergoing compression become less variable in D-period and the inverse for those undergoing tension.

**Extended Data Figure 4.**
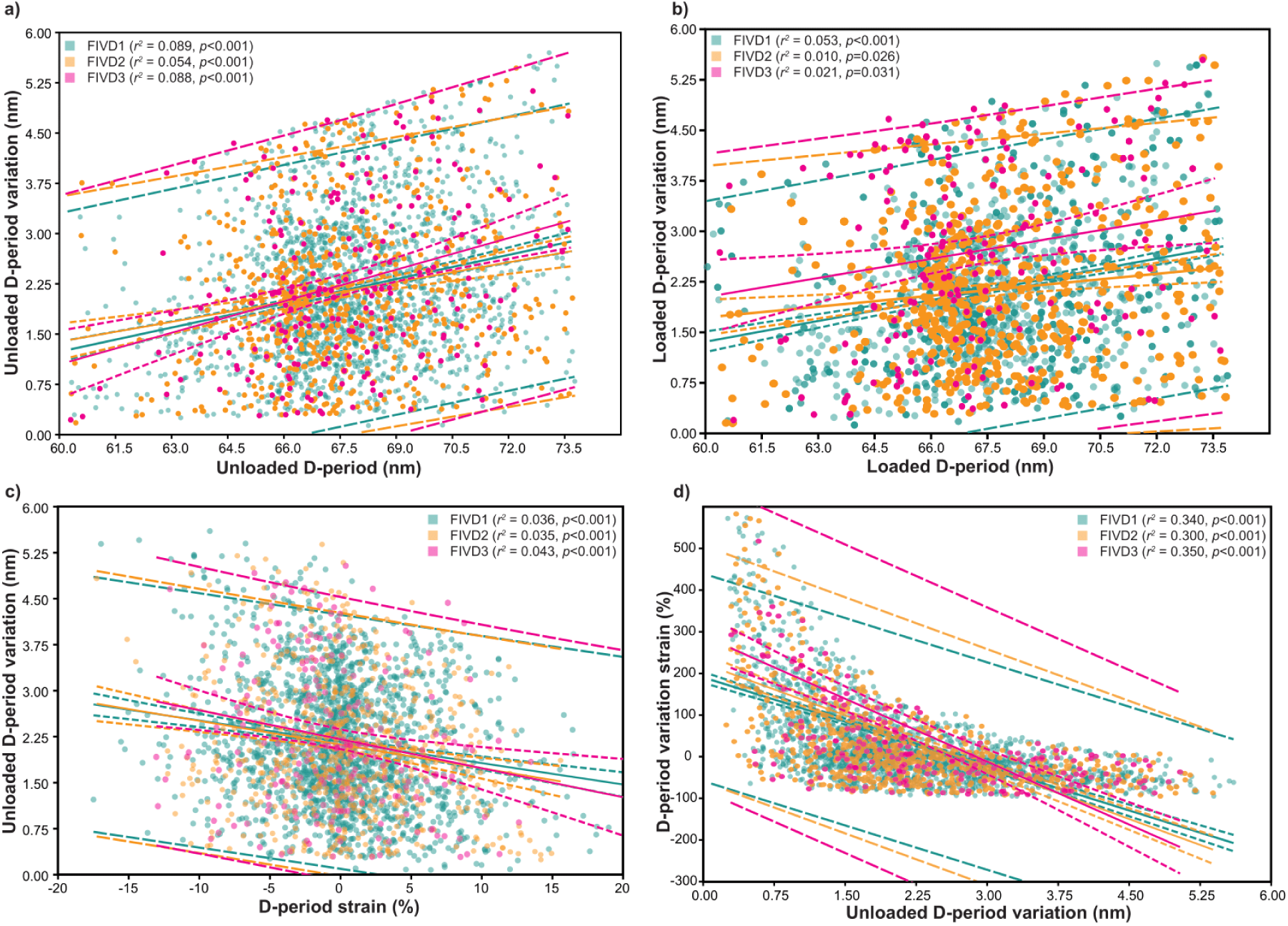
Statistical relationships between mean per-fibre D-period and variation in per fibre D-period for 3 complete IVD samples (Supplementary Table 1).

### Nanoscale fibril properties exhibit lamellar texturing in the AF

Global and spatial patterns in structural and mechanical properties at the fibril to tissue level can be analyzed by the spatial registration of *TomoSAXS* datasets. Mean fibril diameter exhibits two-fold or larger variability across the tissue, with a clear lamellar texture (Fig. 3ei). Closer inspection of fibril diameter distribution (Extended Data Fig. 5h) reveals multiple peaks on a base harmonic of approximately 20 nm. This aligns with prior electron microscopy studies^31^ revealing Type II collagen fibrils form different super-coiled structures from a building block of repeated ∼20 nm micro-fibrils. Combined with the texture seen in Fig. 3ei, these results suggest that super-coiling levels may be different in different regions of the AF and between lamellae.

Algorithmic straightening of maps^32^ for fibrillar-level properties permits comparison of their spatial patterns across individual transects through the AF via line-profiles, providing quantitative support for correlations qualitatively visualized between fibril characteristics and their lamellar texture (Fig. 4 a). Notably, comparisons of D-period pre-strain and fibre orientation (*α*) across shared transects through the posterior AF suggest that pre-strain is broadly conserved within individual lamellae, and neighboring lamellae show contrasting pre-strains suggestive of complementary fibrillar pre-stress (Fig. 4a).

### *TomoSAXS* enables greater sensitivity to spatial variation in nanoscale properties

Statistically, mean values and load-induced changes of each nanoscale property conform to previously reported values in other collagenous soft tissues^8,33^ (Extended Data Fig. 5). However, the range of nanoscale strains significantly exceeds previous estimates from 2D measurements through analogous soft tissues^8,33^ (Extended Data Fig. 5). We suggest that these discrepancies are due to previous studies being restricted to spatial averaging of 2D scattering measurements through tissues and an inability to estimate per-fibre strain. The ability of *TomoSAXS* to reconstruct properties of individual fibres, through isolation of discrete regions of scattering space, allows greater sensitivity to heterogeneities in both their angular value and their scattering that may be under-estimated when averaging across their combined signals (Extended Data Fig. 6). The method thus enables greater understanding of the physiological state of collagenous tissues and reaction to load and can potentially detect local increased fibrillar strains at degenerated lamellae or injured interfaces, which would be lost in spatially-averaged measurements.

The accuracy of *TomoSAXS* for estimating nanoscale parameters has been tested using digital phantoms (Extended Data Fig. 7). We have created a series of digital IVD phantoms by providing indexed fibre geometries from real samples and applying synthetic nanoscale fibrillar properties (Extended Data Fig.7ai-ii). By applying noise in measured background scattering intensity from real SAXS frames (Extended Data Fig.7aiii,av) to simulated signals under analysis (Extended Data Fig.7aiv-vi), we have estimated how accurately the *TomoSAXS* reconstruction reproduces the known parameters. We find significance in both *p* (<0.001) and *r^2^* (0.641-0.894) values for each simulated sample between reconstructed and ground-truth data (Extended Data 7b), which is strongly supportive of our results. Accuracy, measured as mean absolute error (MAER, Methods), was 1.032 nm for mean D-period, 0.280 nm for D-period variation, and 8.532 nm for fibril diameter. Precision, measured as standard deviation of absolute error (SDER, Methods), was 1.37 nm for mean D-period, 0.34 nm for D-period variation, and 15.32 nm for fibril diameter. Comparison of mapped known versus estimated properties also suggest that spatial patterns in known properties are reconstructed (Extended Data Fig. 7b-d).

**Extended Data Figure 5.**
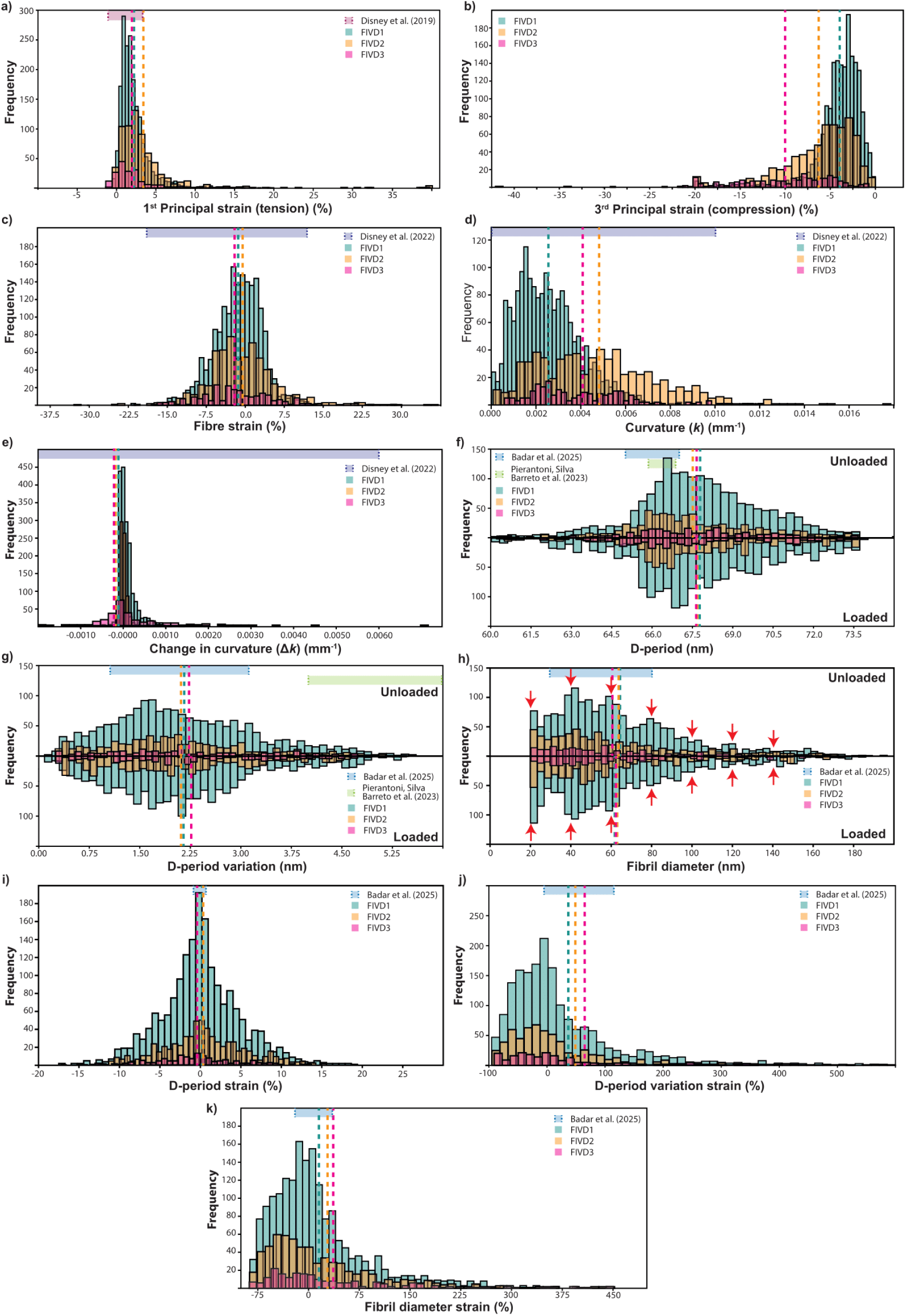
Histograms displaying variation in estimated structural and mechanical properties for samples studied, and comparison to published values.

**Extended Data Figure 6.**
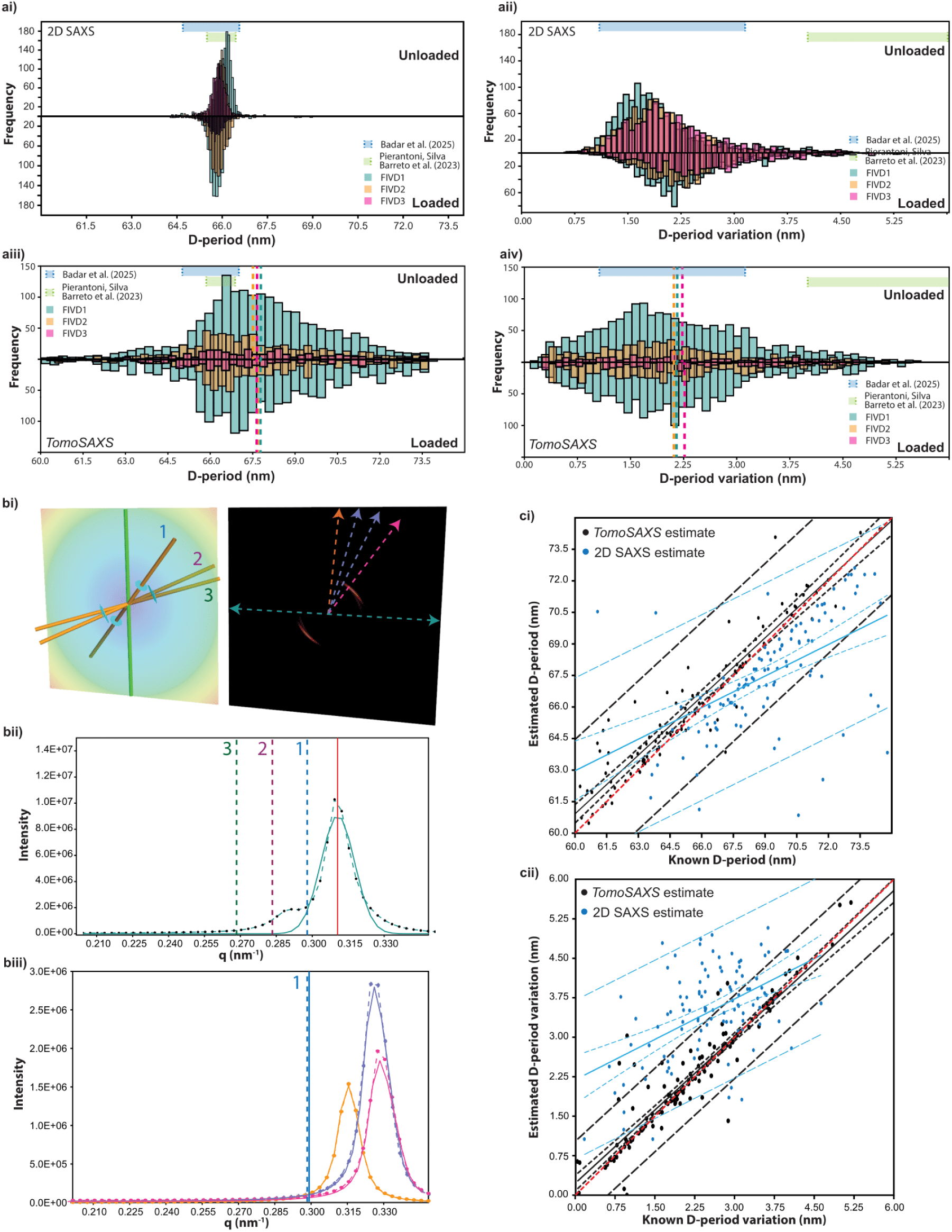
Comparison between performance using *TomoSAXS* versus traditional 2D SAXS approaches for studying nanoscale structure in IVD. a) Comparison between 2D SAXS (above) and *TomoSAXS* (below) analyses of IVD samples. b) Comparison between 2D SAXS and *TomoSAXS* analysis of single voxels modelled using 3D diffraction models, consisting of 3 fibrils of known scattering properties taken from a random distribution with the reported range from real IVD samples. (bi) Estimation of single *q_0_=6π/D* (peak position) for combined scatter through the voxel through fitting a Gaussian model to combined scattering across *q* (red line), compared with known *q_0_* for each fibre. bii) *TomoSAXS* estimation for a single fibre comparing estimated *q_0_* (blue solid line) with known *q_0_* (blue dashed line). c) Comparison between known and fitted D-period (ci) and D-period variation (cii) for 100 simulated fibres using *TomoSAXS* (black) and 2D SAXS (blue).

**Extended Data Figure 7.**
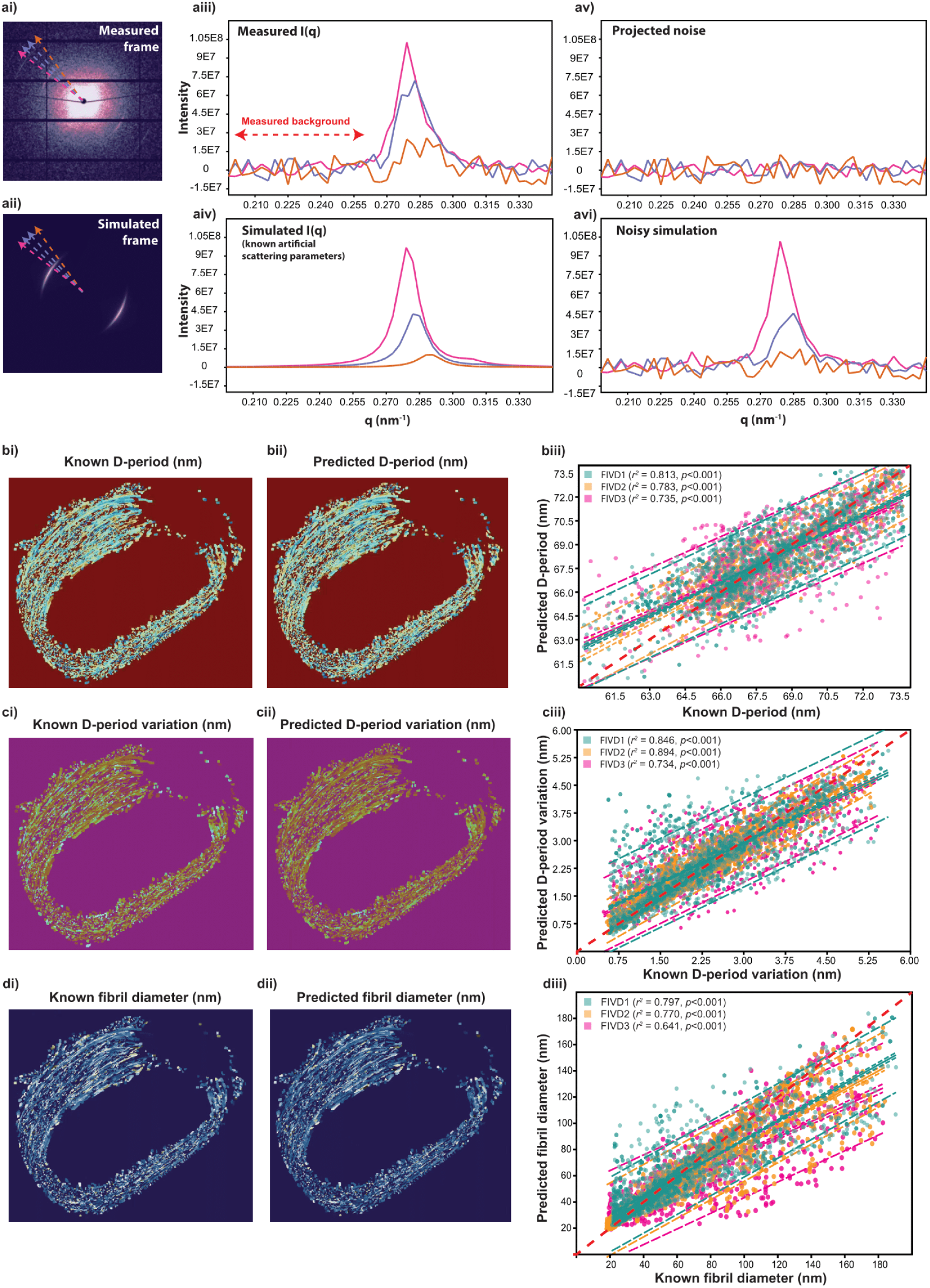
Application of digital phantom data to sample geometries for estimation of accuracy and precision of the *TomoSAXS* technique. a) Schematic illustrating simulation of scattering in a measured frame using known, artificial scattering parameters (retaining fibre index, orientation and estimated scattering amplitude) and application of noise from measured frame. b) Maps and linear regression plots displaying the spatial and statistical accuracy of reconstructions for each nanoscale parameter analysed in *TomoSAXS*. See Supplementary Note 18 for further discussion and presentation of alternative phantoms. Comparison between performance using *TomoSAXS* versus traditional 2D SAXS approaches for studying nanoscale structure in IVD.

### *TomoSAXS* is feasible in intact human organs

We have demonstrated the potential of the *TomoSAXS* technique on human samples, by performing correlative Hierarchical Phase-Contrast Tomography (HiP-CT)^34^ (4-10 µm voxel size) and high energy tomographic SAXS (100 µm voxel size) on whole IVD (∼5 cm width) at ESRF - The European Synchrotron. Challenges for *TomoSAXS* analysis of human IVD arise from the order of magnitude increase in size compared to small animal models, with 1.5 Tb raw HiP-CT data and 800 Gb raw SAXS data generated per sample, compared to 42 Gb pCT data and ∼100 Gb SAXS data for the small animal samples tested here. Preliminary analysis suggests that automated fibre tracing is possible throughout the entire AF (Fig. 4di,ei), and SAXS data provides clear texturing of 3^rd^-9^th^ order meridional collagen peaks (Fig. 4dii,eii-iii). Future applications would include understanding the matrix drivers of healthy and pathological IVD ageing in humans, along with improvements in computational efficiency and speed for processing large datasets associated with intact human IVDs.

## Discussion and outlook

The *TomoSAXS* methodology provides an unparalleled view of macro-to-nanoscale mechanics in the IVD (Fig. 4c). By embedding SAXS data within the structural and mechanical context from pCT and DVC, it enables analysis of nanoscale–tissue interactions within the IVD’s hierarchical assembly. Tissue-level strains show complex patterns, likely linked to non-collagenous AF components such as elastin, thought to enable recoiling of the AF after deformation^16^. Fibre level strains showed a strong correlation with fibre curvature, with highly curved fibres in the inner posterior and posterior-lateral AF reaching up to 20 % strain.

Fibrillar mechanics critically inform strain transmission across scales. Fibrillar pre-strains (initial D-period) and load-induced strain showed strong spatial relationships with fibre-level strains. Their absolute magnitudes suggest most fibre strain is transferred directly to fibrils, with little inter-fibrillar shear, unlike tendons under tension^35^. The averaging effect of commonly used 2D diffraction techniques for studying nanoscale strains and the convolution of scattering from fibres at differing orientations (Extended Data Figure 6) may also contribute to the wider spread of fibril strains seen (up to ±10-15%), which are consistent with tensile tests of individual fibrils^36^.

A strong negative correlation between initial D-period and D-period strain suggests pre-strained fibrils converge toward a neutral state under load. The relationship between mean and variance in per-fibre D-period supports a helical coil model of fibrillar mechanics, loosening/tightening under load (Fig. 4c), akin to prior twisted or coiled fibre models^29,37,38^. This suggests an increase in the variability of stress along fibres as they deviate from a purely compressive (lowest possible D-period) state (Fig. 4c). In this scenario, loading causes increased local swelling pressure to a fibril in a low D-period state, leading to localized regions of high tension (and higher D-period), a phenomenon which is amplified as overall osmotic pressure and residual strain increases. Loss of residual strain in the IVD leads to aberrant cellular mechano-sensing and degeneration^39^; however, methods to measure fibrillar level pre-strain in intact organs have been lacking until now.

The multiscale mechanisms of load response reported here may be incorporated into finite-element modelling of IVD biomechanics^40^ and explain complex non-linear load responses. Collagen fibrils store stress and strain energy through osmotic pressure shifts, generating hundreds of MPa during dehydration^41^. These correlations have potential clinical relevance in musculoskeletal degeneration, with reduction in swelling pressure due to synthetic aging shown to result in a reduction of fibrillar pre-strain in articular cartilage^42^. By spatially deconvolving 3D pre-strain maps (Fig. 3), *TomoSAXS* surpasses prior 2D SAXS studies that reported only depth-averaged changes. Indeed, due to the range of fibre-orientations and variable tissue-level strain in IVD, conventional SAXS would have reported an average fibril-strain of zero, missing the rich intra-tissue texture and multiscale correlations. Other soft tissues like arteries exhibit a multi-layered structure with variable fibre orientation and internal pre-stress; changes in ultrastructural pre-strain at the fibrillar level may also play important biomechanical roles in such systems^43,44^.

**Extended Data Figure 8.**
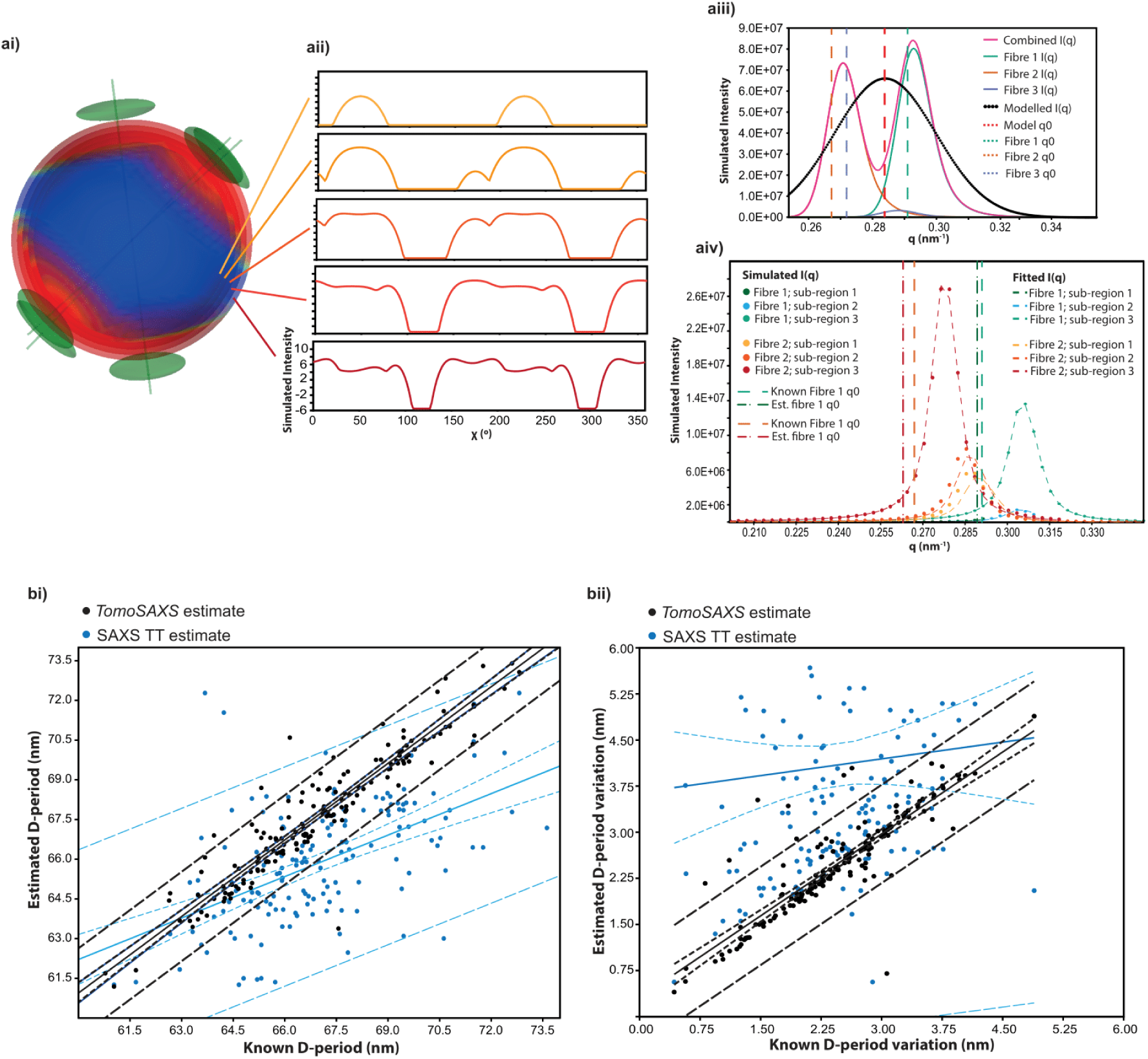
Comparison between SAXS tensor tomography (SAXS-TT) and *TomoSAXS* for reconstructing scattering parameters of individual fibres. a) Schematic illustration comparing the SAXS-TT and *TomoSAXS* methods for estimating *q0* for three fibres in a single voxel of known scattering parameters. ai) Three-dimensional (3D) visualisation of 3D reciprocal scattering for each fibre based on their angular orientation and scattering parameters. Intensity stratified into 5 consecutive “shells” sampling reciprocal space along *q*. I(*χ*) plots for these shells are provided in (aii). aiii) estimation of *q0* (red dashed line) via fitting Gaussian functions (black dotted line) to amplitude values for model fits to 200 shells between *q*’s of 0.255-0.35, and comparison with known *q0* for each fibre. aiv) Individual *TomoSAXS* fits and *q0* estimation for fibres that provided sufficient independent scattering. b) Comparison between known and fitted D-period (bi) and D-period variation (bii) for 100 simulated fibres using *TomoSAXS* (black) and SAXS-TT (blue).

*TomoSAXS* thus presents a powerful addition to multimodal X-ray imaging for spatial surveys of nanoscale properties^45^, offering key advantages over approaches such as SAXS tensor tomography^19^ (SAXS-TT) for the study of fibrous materials. By spatially registering SAXS data to collagen fibres defined by their 3D orientations (from pCT), *TomoSAXS* enables fibre-specific — rather than voxel-specific^19^ — assessment of nanostructure. This allows direct analysis of heterogeneous fibre scattering profiles, with resolution determined by pCT (1.6 µm voxel size here) rather than the SAXS beam size (20 µm here, Extended Data Fig. 8), and estimation of fibre-specific nanoscale strain. The combined CT-DVC-SAXS approach is essential to resolve fibril-level strains, with DVC also providing strain information at the fibre and tissue scales. Tracking fibres across slices allows interpolation methods that preserve fibre continuity. Using the structural model of the fibril scattering function, rather than the more general spherical harmonic approach, allows more accurate measurements, albeit currently at the cost of increased computational time and greater specificity of the scattering functions for the material investigated.

Scan time remains a limitation, particularly for the investigation of mechanics in viscoelastic samples, as samples must be stable during scanning. The need for stress-relaxation means that the dynamic response of materials immediately following loading is not captured. Fast (5 s) pCT scan times have been used to capture dynamic loading of cartilage^46^, and could be combined with DVC to determine tissue and fibre level strains during loading. Radiation dose is also critical, though controlled exposures here (7.3 kGy SAXS; 27 kGy pCT) have been shown to not affect mechanics in these samples^22^ and other soft collagenous tissues^46^. Extending the application of *TomoSAXS* to mineralised tissues such as bone must address diffuse scattering from the mineral phase and the difficulty of resolving collagen orientation from pCT.

Looking ahead, the *TomoSAXS* methodology is broadly applicable to other collagenous and soft biological materials. We have demonstrated the feasibility of performing *TomoSAXS* on intact human IVD, with potential to extend the technique to other organs including the heart and lungs. Whilst synchrotron radiation was used here, *TomoSAXS* could also be undertaken with laboratory X-ray sources at the expense of markedly increased acquisition times. The measurement of fibre orientations from pCT could be replaced with other 3D microscale imaging techniques, such as optical coherence tomography^47^. Extending to engineered materials, our 3D diffraction modelling software may also be developed for other periodic and angularly dependent materials including nanostructured polymers^48^ and carbon fibre composites^49^.

## Methods

### Sample acquisition and preparation

Samples originated from male 8-week-old Sprague Dawley rats kept in controlled conditions (12:12 hour light:dark cycles, 22 °C and ad libitum access to food and water) at the University of Manchester Biomedical Sciences Facility. All tissue collection procedures were performed in accordance with the UK Animals (Scientific Procedures) Act 1986, local regulations set by the UK Home Office with institutional approval obtained from the University of Manchester Animal Welfare and Ethics Review Board under the Home Office Licence (#70/8858 or I045CA465). Whole spines were dissected and snap frozen in liquid nitrogen prior to storage at -80° C. Thawing and further dissection were performed within 24 hours of experimentation, as described by Parmenter et al^22^. This resulted in whole IVD samples (from vertebral levels thoracic 10 to lumbar 2, Supplementary Table 1) with half vertebrae on either side.

Samples were mounted in resin-printed holders (Supplementary Fig. 1) using cyanoacrylate glue (VetBond; 3M Science). A Kapton polyimide tube enclosed each specimen, leaving ∼0.5 mm exposed. Holders were placed in a Deben CT500 micromechanical tester (100 N loadcell, ±1% accuracy; Deben, UK) with a custom open-frame design^22^ (Supplementary Fig. 1). Displacement was recorded using a 10 mm extensometer (3 μm resolution, 1% linearity). The open-frame configuration facilitated alignment and hydration of samples while minimizing beam-path obstruction during SAXS. Sample hydration was maintained through immersion in phosphate-buffered saline (PBS), supplied via a capillary at the holder base from a 100 ml syringe through a stopped-flow module (Bio-Logic, France). A continuous flow preserved fluid level despite minor leakage from the 3D-printed holder.

### Overview of SAXS/pCT experiment

While simultaneous SAXS and pCT would have been ideal, in the absence of a dedicated SAXS/pCT beamline at Diamond Light Source (DLS, Harwell, UK), a sequential approach was taken (SAXS(I22) preloaded →sCT(I13) preloaded →sCT(I13) loaded →SAXS(I22) loaded), as illustrated in Extended Data Fig. 1, with special stabilized sample holders and identical mechanical rigs at both beamlines to ensure comparability (Supplementary Fig. 1).

### Acquisition of tomographic SAXS data

SAXS data were acquired at the I22 beamline of the DLS synchrotron (Experiment SM29784-8, 12–15 December 2023). A Deben CT500 was mounted to a rotation stage, on top of the sample table. A Dectris Pilatus P3-2M detector was placed at a distance of 9 m from the stage, with a vacuum tube in between to minimise air scatter. All SAXS measurements used a beam energy of 20.2 keV and beam size of 20 µm. Each sample was first mapped at 0° angular orientation using a raster map of approximately 100*100 SAXS frames using a 50 ms exposure time, and a 20 µm step size. Maps were inspected to identify the IVD, and a 5*1 mm region of interest was assigned within the IVD for tomographic SAXS scans. SAXS scans were undertaken over raster maps of approximately 250×10 SAXS frames with a step size of 20 µm using 150 ms exposure time for 9 equally spaced angular orientations between -85° and 95° (to minimize blocking by mechanical testing support rods). Radiation dose for a full single axis SAXS tomography was 7.3 kGy (Supplementary Note 1).

### Acquisition and reconstruction of pCT data

pCT was conducted at the Diamond-Manchester Imaging Branchline I13-2, DLS (Experiment SM29784-9, 12–14 December 2023). The setup followed established protocols^22^ (mean energy 27.6 keV, 1800 projections, 0.5 m sample–detector distance), with only the exposure time adjusted to 0.15 s to account for a reduced ring current of 200 mA due to a radiofrequency cavity issue. The insertion device gap was set to 5 mm during scanning and widened to 7 mm for alignment and low-resolution scans to reduce dose. Initial scans used a 1.25× objective (2.6 µm voxel size, 6.7 × 5.6 mm field of view) to capture the sample environment, including the Kapton tube for SAXS co-registration. Subsequent scans employed a 2× objective (1.625 µm voxel size, 4.2 × 3.5 mm field of view) before and after compression. Radiation doses were 1.35 kGy (2.6 µm voxel size) and 27 kGy (1.625 µm voxel size) per scan (Supplementary Note 2). Tomograms were reconstructed as described previously^22^.

### *In situ* mechanical testing

Samples were held under axial compression, applied in the laboratory vertical z-direction, during scanning. For a reproducible mechanical protocol, two identical Deben CT500 mechanical testing rigs with 100 N load cells were used during the experiment, one placed at beamline I22 (provided by Imperial College London) and the other placed at beamline I13-2 (provided by the University of Manchester at Harwell). A preload of 1 N, corresponding to approximately 40 % the animals body weight (260 ± 33 g), was applied at I22 prior to SAXS scanning (Extended Data Fig. 1b). This preload generated an intradiscal pressure of ∼0.1 MPa (∼10 mm^2^ cross-sectional area), comparable to pressures measured in human subjects lying prone^50^. Samples were allowed 20 minutes stress relaxation before scanning to minimise motion. Following SAXS, samples were clamped in position and transferred to I13-2 for pCT (Extended Data Fig. 1b).

Clamped samples were placed in the mechanical testing rig at I13-2 and secured at the top (Extended Data Fig. 1c). The bottom platen of the rig was slowly raised until just touching the clamped sample holder, which was then secured to the rig. The clamps were removed from the sample holder prior to preload sCT scans. After the first sCT scan at 1.6 micron voxel size, a 50 micron displacement was applied to the sample, leading to a 2-5 % reduction in IVD height (Supplementary Table 1). Again, samples were given ≥20 minutes stress relaxation before scanning. After imaging, the sample was clamped in its loaded state before being removed from the rig and transported back to I22 (Extended Data Fig. 1c). At I22 the sample was placed back in the mechanical testing rig and clamps removed before taking the loaded SAXS scans, which used the same protocol as described in the section “Acquisition of tomographic SAXS data” (Extended Data Fig. 1d).

### Fibre tracing, orientation calculation and DVC analysis of sCT data

Fibre tracing of the preload 1.6 µm images was performed using the Avizo XFiber module (Avizo 3D 2023.2), with parameters used for tracing given in Supplementary Note 3. After fibre tracing, fibres clearly corresponding to imaging artefacts – such as streaks from mineralised tissues and ring artefacts from reconstruction – were removed from further analysis. Fibre spatial graph data was exported as a .xml file, and fibre identification number conserved throughout all further analysis.

Fibre orientation was calculated using a custom written Matlab script (Matlab R2023a, Supplementary Note 4), resulting in fibre ID, x, y, z, α and β orientation values for points placed every 6.5 µm along each fibre. Fibre ID, α, and β values were converted to 3D volume images for input into the *TomoSAXS* reconstruction (Supplementary Note 5).

A local DVC analysis with flexible point cloud specification was performed using the open-source software iDVC (https://tomographicimaging.github.io/iDVC/)^9,51–54^. The fibre-based coordinates generated above were used as the DVC point cloud defining the locations of sub-volumes used for tracking (Fig. 2e), resulting in 0.7 – 1.4 million DVC measurement points/ sub-volumes for each sample (Supplementary Table 1). Spherical sub-volumes with a diameter of 40 voxels (65 µm) were matched between the preload and loaded images using a zero-normalised sum-of-square difference function for optimisation and 12 degrees of freedom (translation, rotation, and 1^st^ order strain, Fig. 2g). 6000 sampling points were placed within each sub-volume for matching. The displacement of the first sub-volume in the point cloud was defined by manually tracking a landmark between the preload and loaded images using ImageJ.

Two strain measurement techniques were applied to the displacement fields measured from DVC, corresponding to different length-scales probed. For tissue-level strains, the differential of a polynomial fit to the displacement field was used to calculate the Lagrangian strain tensor at each point^53^. The 75 closest points to the measurement point were used for fitting to smooth out noise in the displacement field and improve accuracy. 1^st^ principal strain was derived from the maximum positive eigenvalue and 3^rd^ principal strain the maximum negative eigenvalue of the strain tensor. The direction of the corresponding eigenvectors relative to the local fibre direction was calculated as an angle between 0° (parallel to fibre direction) and 90° (orthogonal to fibre direction, Extended Data Fig. 3g). Radial and circumferential strains were calculated by projecting the strain tensor along the local radial and circumferential directions (Extended Data Fig. 3a, Supplementary Note 6). Fibre strains were calculated using a custom written Matlab script (Matlab 2023a), which performs a polynomial fit to displacement along the fibre direction (Supplementary Note 7, Supplementary Figure 2).

The double projection zero strain method^22,52^ was used to quantify the accuracy of DVC displacements and strains (Supplementary Note 9). Zero-strain displacement accuracy (mean absolute error) was 99.4 nm and precision (standard deviation of absolute error) 148 nm. Tissue zero-strain accuracy was 0.17 % and precision 0.23 %. Fibre zero-strain accuracy was 0.07 % and precision 0.15 %. DVC was able to track a virtual deformation of 10 % compression with a displacement accuracy of 198 nm, tissue strain accuracy of 0.40 %, and fibre strain accuracy of 0.45 %.

To calculate orientation for the same fibres in each sample after loading, DVC displacements were applied to the fibre points in Matlab (Supplementary Note 10). Orientation of the deformed fibres was calculated, and loaded Fibre ID, α, and β 3D volume images generated for input into the *TomoSAXS* reconstruction.

### 3D diffraction modelling

3D diffraction modelling was applied by simulating the meridional X-ray diffraction peaks of collagen fibrils with nanoscale fibre symmetry, under user-defined experimental conditions (wavelength, lattice spacings, angular tilts), and accounting for the variable intersection of the 3D scattering intensity with the Ewald sphere. Implementation was done in a custom Python library developed by us and is a more complex version of the model presented by Badar et al^8^. Briefly, the meridional scattering arising from the D-periodicity is simulated by a set of 3D semi-flat ellipsoidal functions, whose location in 3D space is at multiple harmonics of the inverse D-period (q_0n_ = 2πn/D), radial width in *q* (w_a_) is related to fibril pre-strain heterogeneity, and angular width (w_μ_) is related to inverse fibril diameter and intrafibrillar tilting, parameterized by a degree of circularity of the ellipsoid function (δ; 0 = straight function; 1 = circular ellipsoid). The need for a semi-flat, rather than completely flat, meridional ellipsoid came from the experimental observation that even for tissues with uniaxial fibril orientation, the meridional ellipse showed some arcing. We believe this may be related to intrafibrillar tilt of the microfibrils^29,55^.

Due to the intersection with the Ewald sphere (radius ∝ 1/λ), the intensity and pattern of the resulting 2D fibril diffraction depends on user-input 3D tilt angles α, β of the fibril, which will be a crucial factor in deciding which fibres contribute to scattering, as well as the beam wavelength. The functions for fibril scattering can be calculated at any point across the diffraction condition of the sphere both in 2D (Fig. 2bi), as well as in 1D (e.g. azimuthal (χ) and radial (q) plots) (Fig. 2c) by averaging in the complementary axis.

### Overview of *TomoSAXS* algorithm

Here, the foregoing library, in conjunction with the sCT/DVC data (for α, β) was used to reconstruct the third order collagen diffraction peak (which typically has intensity between wavevectors ∼0.26-0.32 nm^-1^). The *TomoSAXS* implementation works via a series of discrete modules in the Python (3.10) environment, summarised here.

- **Module 1** spatially co-registers the sCT and SAXS scans, from which the individual collagen fibres in each SAXS beam path at all tomographic rotations can be found.
- **Module 2** performs background correction of the SAXS signals, accounting for PBS, Kapton tube and air scattering, building on the *adcorr* Python package^56^. Importantly, the sCT and co-registered SAXS scans from **Module 1** allows accurate correction using the variable thickness of the sample through each SAXS beam-path by adjusting the proportion of dispersant to sample, compared to assuming a uniform thickness in Pauw et al^56^.
- **Module 3** provides initial estimates for per-fibre SAXS amplitude, using singular value decomposition (SVD) and the 3D diffraction model.
- **Module 4** – the core of the reconstruction – extracts the nanoscale parameters in an iterative way, by scanning through the SAXS tomography along χ- and rotation axes to find angular regions where either a single fibre, a pair of fibres or triplet of fibres dominates the observed scattering over a given angular sector (10° in width). These cases are fitted to the diffraction model to give mean D-period (inversely to peak centre; Equation 1); variation in D-period (peak width along q); and fibril radius (inversely related to peak width along χ). The process continues until all solvable fibres have been fitted.
- **Module 5** spatially reconstructs and outputs the estimated metrics, and estimates fibril-level (nanoscale) strain in biomechanical loading, using DVC estimates of per-fibre (microscale) deformation to locate the same fibre in the unloaded and loaded datasets.

### Module 1: Spatial co-registration of fibre orientation data

Spatial co-registration of SAXS and sCT datasets was achieved through identifying fiducial markers (like the mineralized vertebral endplates) in the samples which gave distinct signals in SAXS and pCT. The Kapton tube of the sample environment was used for horizontal alignment and the mineralised tissues of the vertebral endplates used for vertical alignment. High resolution (1.625 µm) pCT images did not capture enough Kapton tube for co-registration, so were spatially registered to low-resolution (2.6 µm) images. Both sets of images were down sampled to 6.5 µm voxel size and the offset between them calculated in x, y, and z coordinates. Fibre orientation tiff stacks (absolute voxel size 8.125 µm) were padded to the size of the original high resolution (1.625 µm) reconstruction (512*512*432 voxels) and centred using the offset between high- and low-resolution images, so that all pCT data was spatially aligned.

Vertical registration between SAXS and pCT involved identifying the border of the mineralised vertebral endplates in the SAXS intensity map generated during coarse mapping at 0°, and in the 2D total phase contrast intensity maps of a sample along the same angular orientation from pCT data, and calculating the vertical offset between them. Horizontal registration was achieved by identifying the Kapton tube from its characteristic peaks in SAXS intensity, and manual segmentation of the tube from pCT data. For each SAXS tomography orientation (*n* = 9, -90° to 90°), the fibre orientation, index, and Kapton data for vertically isolated SAXS slices was rotated to the corresponding angle (using a custom function for maintaining fibre data integrity during rotation; Supplementary Note 13). The location of the left-hand edge of the Kapton tube was then compared and aligned between SAXS and pCT data. This allowed finding the indexed fibres that contributed to SAXS scattering along each beam-path of the raster scan in the specific SAXS slice. For each frame, the orientation, index, and number of CT voxels comprised by each indexed fibre along its beam path was saved, along with the calculated distance between each fibre and the centre of the beam path, which was used as a weighting factor for model fitting. This data was saved as individual matrices, along with calculated total specimen thickness (along beam path) for each frame.

### Module 2: Background correction

Background correction was undertaken for each individual SAXS frame following the method outlined in Pauw et al^56^. A separate scan using the same parameters as the SAXS tomography was undertaken for a single slice (a slice represents a 1D scan x n=9 rotations) through 1) an empty Kapton tube and 2) a Kapton tube filled with PBS, to represent background SAXS signals from both the sample holder (Kapton tube) and the dispersant (PBS). These signals were algorithmically subtracted from each sample SAXS frame using the *adcorr* python library (https://github.com/DiamondLightSource/adcorr), adjusting the ratio between sample and dispersant based on the sample thickness for the respective frame (Module 1). The resultant background corrected data were saved as .h5 files for each TomoSAXS slice.

### Module 3: Per-fibre SAXS amplitude estimation

The first analytical step in the TomoSAXS algorithm makes an initial estimate of the amplitude of SAXS scattering for individual fibres using single value decomposition (SVD), using a mean value for the other peak characteristics like position, radial and angular widths. This is justified by the small variability of the peak position (from 2D data) across different sample locations. The estimation step is used to identify locations of strong scattering for specific fibres enabling fibre-by-fibre solution in the next Module 4 step.

At a given beam path in the sample *x*, and a rotation angle r, the angular intensity of the 3^rd^ order collagen peak q_0_ can be written as

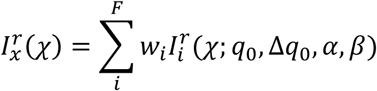

where the sum in *i* is taken over all fibres in the volume, with weighting factors either 0 (if outside of the beam path) or proportional to the intensity at the fibre location (assuming a Gaussian beam profile with known shape). Since all fibres are taken to have the same average peak location and width parameters in this initial estimation step, the contribution from each fibre 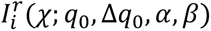 can be calculated up to a scaling factor proportional to the amplitude factor A_i_, including the geometrical transformations needed to account for fibre orientation with respect to the beam. Hence when represented for each *x*, *χ*, *r* the above equation can be expanded into an overdetermined set of linear equations which can be solved by SVD.

A python dictionary was created with entries for each SAXS frame comprising of 0-180° Intensity(χ) (herein I(χ)) azimuthal integrations between 0.26-0.32 nm^-1^ q, binned to 36 averaged points across χ and smoothed using 1-dimensional Savitzky-Golay filtering (window length = 9, polynomial order = 1). Two matrices were then created; the first consisting of a single vector comprising the concatenated I(χ) integrations for all SAXS frames (LHS in the Equation above); the second with columns comprising the contribution of each fibre to the same binned I(χ) intervals predicted for each SAXS frame using 3D diffraction modelling (RHS in the Equation above). The amplitude for each fibre was then estimated by non-negative linear least squares regression, regressing predicted versus measured I(χ) data. A python dictionary was constructed, with entries for each indexed fibre consisting of its index, α and β angles and an empty entry for fitted nanoscale parameters. This dictionary was used to calibrate registration matrices (see above) for each rotation angle of the SAXS tomography, creating a series of other dictionaries.

### Module 4: TomoSAXS fitting

#### Initial intensity simulation

As shown in Fig. 2f-h, *TomoSAXS* reconstruction is based on fitting fibre diffraction models of intensity to measured I(q) profiles across the angular ranges (Fig. 2g) of independent scattering signals from either i) a single fibre, ii) a pair or iii) a triplet of overlapping fibres (Supplementary Table 2). We first calculate simulated angular SAXS intensity profiles using estimated amplitudes from Module 4. A Python fibre dictionary was constructed, containing its index, α and β angles, initial estimated scattering amplitude, and an dictionary for fitted nanoscale parameters. The dictionary was referenced against registration matrices (see above, Module 1) for each rotation angle of the SAXS tomography, creating a series of individual orientation-specific dictionaries, indexed by beam path.

For each beam-path in the orientation-specific dictionary, a dictionary indexed by fibre index stored information for every fibre in the beam path (including initial estimated amplitude), with a weighting term from the calculated distance of the fibre to the beam-centre (Supplementary Note 13) where *d* is the distance from beam centre and *w* the beam diameter:

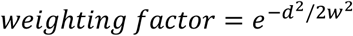

These dictionaries were used to create simulations of the angular scattering intensity of the 3^rd^ order meridional peak of every beam-path using 3D diffraction modelling, using tilt angles obtained from pCT and the averaged nanoscale peak metrics (q_0_ = 0.2856 nm^-1^; w_a_ = 0.005 nm^-1^; w_µ_ = 0.2 nm^-1^; δ = 0.5 see Supplementary Note 14 for rationale).

#### Fitting single, double and triplets of fibres

The angular intensity profile was scanned to iteratively solve individual or small groups of fibres, rather than fitting all fibres to the data, simultaneously.

##### Single fibre fit

For each fibre *k* in a beam path *p* at a rotation *r*, its individual angular intensity contribution 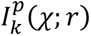 to the total intensity *I*^*p*^(*χ*; *r*) was calculated. If the following condition was fulfilled for an angular range 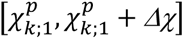 between 0 and 2π (Δχ=10°), and a single fibre dominance threshold *t_s_*=0.90:

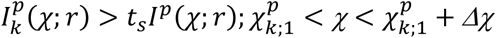

then the fibre *k* was identified as being the dominant contribution to the SAXS signal in the 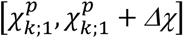 angular range. The index, beam-path location, and angularly independent sector 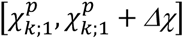 was recorded for all fibres that met the user-defined criteria for independent scattering were stored in a Python list.

Next, these fibres were fitted to extract the nanoscale (fibrillar) peak metrics, by calculating *n*_*χ*,*c*_ (default = 3) equally spaced slices across the 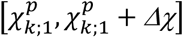 angular region, with a width Δ*χ*_*c*_ (default = 6°). These were simultaneously fitted with 3D diffraction models for the fibre, using the minimize() function of the *lmfit* Python library (https://lmfit.github.io/lmfit-py/fitting.html) and the Nelder-Mead algorithm. The angles α and β and the weighting factor of the fibre were fixed parameters, and q_0_, w_a_, w_μ_, amplitude, and δ were set as fit parameters (see Supplementary Table 2 for settings and Supplementary Note 15 for rationale). If the fits passed several quality thresholds, both relating to absolute values of the parameters and statistical quality of fit (standard error: threshold = 25%), the fit values were added to the dictionary entry for the respective fibre. Instances where the fit to the fibre did not meet quality thresholds were saved and added to a separate python list allowing them to be discarded from future investigation. This process was repeated for all the fibres in the list.

Since successful fits allowed more potential single-fibre instances to be identified (already fitted fibres could be deconvolved from the scattering in all beam-paths where they occur), the process was repeated until no more instances of independent scatter for single fibres could be found for any beam paths in the SAXS tomography.

The above process isolated and fitted the instances with the strongest independent scattering signal (∼10-20% of indexed fibres per-slice), providing robust fits for further deconvolution and fitting of other fibres occurring in the next step of the reconstruction algorithm.

##### Overlapping fibre fit

In the next step, the reconstruction algorithm included fitting of instances where two to three fibres produced a combined scattering that meets the thresholds for independent scatter. E.g. for all pairs of fibres *k*_1_, *k*_2_ in the beam-path *p*, we checked whether the following condition was fulfilled for any 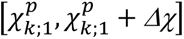 and a double fibre dominance threshold *t_d_*=0.95:

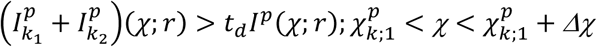

For multiple fibre fits, validation simulations (Supplementary note 15) have shown that the dominance of one fibre’s scattering intensity can bias the fit of the other fibre(s) and lead to inaccuracies in fits. Hence, upon identification of a pair of fibres *k*_1_, *k*_2_, the relative combination of scatter from each fibre was estimated. Only instances where each fibre represented a minimum of 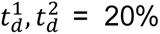 of the total scatter across *Δχ*_*d*_ > (0.90) of 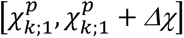 were considered for fitting. Instances that passed this test were then fitted using the minimize() function as above.

After each fit, the parameter database was updated, and each angular intensity profile was scanned to identify any new solvable fibres (single, double or triple) which could be fitted by deconvolving already fitted fibres.

The foregoing steps were looped through all beam paths at all rotations, for a user-defined maximum number *n*_*l*_ = 10 times, recording the ratio of fitted versus total indexed fibres at the end of each loop. If no increase in fitted fibres was observed at the end of a loop, the loop was terminated as all possible fibres had been fitted.

The entire *TomoSAXS* pipeline was conducted on a per-slice basis (per-horizontal 20 µm SAXS tomography slice) using cluster computing (specifically the DLS computing cluster). This allowed each slice to be run in parallel to maximise efficiency. The complete process across each of the five modules was completed within six-seven days, with each module triggered individually in an iterative manner. *TomoSAXS* fitting (four-five days) resulted in a mean proportion of fitted fibres of 60.08% per-slice. This included multiple fits for differing instances of individual fibres found within different *TomoSAXS* slices. It is important to note that this does not correspond to the proportion of the total sample represented in each slice being fitted, as *TomoSAXS* operates on a per-fibre basis as opposed to a per-voxel basis (as in SAXS tensor tomography^19^). Reconstructions are spatially heterogeneous and dependent on the density and orientation of fibres within the respective slice.

### Module 5: TomoSAXS reconstruction

Once fitting was complete, fitted parameters were spatially reconstructed for scanned fibres to create volumetric maps of per-fibre values. This was performed per-slice for each parameter, by creating blank maps of the same shape as fibre index maps (created during the registration process; Supplementary Note 13) and assigning metric values to the voxels assigned to the index of every solved fibre in index maps. For fibres with fitted values in multiple *TomoSAXS* slices, if slices remained unfitted between these points, they were provided with values between fitted values estimated using linear interpolation.

#### Per-fibre nanoscale strain reconstruction

To estimate strain in nanoscale parameters under compressive load, indexed fibres were isolated that provided fits in both unloaded and loaded data. For each unloaded fit, the estimated vertical deformation of this point (and thus position in the loaded *TomoSAXS* scan) was estimated using the vertical (*z* axis) deformation measurements provided by DVC analysis (Supplementary Note 17). Strain in each parameter was calculated as the percentage difference between unloaded and loaded *TomoSAXS* fits for each point.

### Validation using digital phantoms

The spatial calibration between *s*CT and SAXS data allowed estimation of the accuracy of fits supplied by the *TomoSAXS* technique, by providing known artificial nanoscale parameters to each indexed fibre in each *TomoSAXS* slice (maintaining fibre orientation, initial estimated scattering amplitude, and per-instance weighting, Supplementary Note 18). Experimental noise was added to the simulated SAXS data, and accuracy and precision of nanoscale parameter estimates calculated as MAER and SDER.

### *TomoSAXS* of human IVD

Static imaging of a whole, formalin fixed, human IVD (spinal level thoracic 11-12, 64-year-old male donor, no known spinal pathologies) was performed at the BM18 beamline of the ESRF using hierarchical phase-contrast tomography^34^ (Supplementary Note 21). The entire IVD was captured with 10 µm voxel size, and a local zoom with 4 µm voxel size performed centrally to enable clear visualisation of collagen fibres throughout the complete organ (Fig. 4d,ei). Correlative high energy tomographic SAXS was performed at ESRF beamline ID31 (Supplementary Note 22).

### Statistical Analysis

Ordinary Least Squares (OLS) regression of per-fibre strain (%) versus curvature (*k*): curvature = 0.035 * fibre strain + 0.001; *r^2^* = 0.480, *p* = 0.0001. OLS regression of per-fibre strain (%) versus D-period strain (%): D-period strain = 0.347 * fibre strain + 0.024; *r^2^* = 0.138, *p* = 0.0001. OLS regression of initial D-period (nm) versus D-period strain (%): D-period strain = -1.357 * D-period + 0.040; *r^2^* = 0.312, *p* = 0.0001. OLS regression of initial D-period (nm) versus D-period variation (nm): D-period variation = 0.107 * D-period + 0.01; *r^2^* = 0.089, *p* = 0.0001. OLS regression of initial D-period (nm) versus D-period variation (nm): D-period variation = 0.107 * D-period + 0.01; *r^2^* = 0.089, *p* = 0.0001. OLS regression of D-period strain (%) versus D-period variation (nm): D-period variation = -0.035 * D-period strain + 0.005; *r^2^* = 0.067, *p* = 0.0001. OLS regression of D-period variation (nm) versus D-period variation strain (%): D-period variation strain = -56.89 * D-period variation + 1.891; *r^2^* = 0.340, *p* = 0.0001. OLS regression of known D-period (nm) versus estimated D-period (nm): estimated D-period = 0.646 * known D-period + 0.010; *r^2^* = 0.813, *p* = 0.0001. OLS regression of known D-period variation (nm) versus estimated D-period variation (nm): estimated D-period variation = 0.826 * known D-period variation + 0.0006; *r^2^* = 0.846, *p* = 0.0001. OLS regression of known fibril diameter (nm) versus estimated fibril diameter (nm): estimated fibril diameter = 0.704 * known fibril diameter + 0.007; *r^2^*= 0.797, *p* = 0.0001.

## Acknowledgements

Laboratory space and facilities were provided by the Research Complex at Harwell. We thank S. Marussi for his support with the adaptation of the mechanical testing rig. We acknowledge the University of Manchester at Harwell for providing the mechanical testing rig. We thank A. Sharma, L. Forster, J. Liu, S. Marathe, and Y. Zhou for their help on beamtimes. We acknowledge funding from the UK Engineering and Physical Sciences Research Council (EP/V011235/1, EP/V011006/1, EP/V011383/1, EP/V011065/1), UK Medical Research Council (MR/R025673/1 and MR/V033506/1), Royal Academy of Engineering (CiET 1819/10), Chan Zuckerberg Initiative (CZIF2022-316777), ESRF HOAHub MD-1389, MD-1290, & ME-1735, & Diamond Light Source long-term project (SM29784). A.L.P. acknowledges support from the i4health Centre for Doctoral Training. H.S.G. acknowledges support from BBSRC (BB/R003610/1).

