## Supplementary information for "Multimodal X-ray imaging reveals hierarchical fibre mechanics"

### Supplementary Materials

**Supplementary Table 1 –** Spinal level, disc height, bulk IVD strain, fibre tracing, and DVC values for each of the 3 samples analysed.

| **Sample:** | **IVD 1** | **IVD 2** | **IVD 3** |
|---|---|---|---|
| **Spinal Level** | Lumbar 1 – 2 | Thoracic 11 – 12 | Thoracic 10 – 11 |
| **Mean distance between endplates at preload (mm)** | 1.054 mm | 0.658 mm | 0.692 mm |
| **Minimum distance between endplates at preload** | 0.563 mm (346 voxels) | 0.186 mm (114 voxels) | 0.296 mm (182 voxels) |
| **Mean distance between endplates loaded** | 1.033 mm | 0.631 mm | 0.657 mm |
| **Bulk compressive strain** | 2.00 % | 4.10 % | 5.06 % |
| **Strain at minimum distance between endplates** | 5.21 % | 15.21 % | 11.80 % |
| **Minimum length used for fibre tracing** | 520 µm (320 voxels) | 260 µm (160 voxels) | 162.5 µm (100 voxels) |
| **Number of fibres used for analysis** | 9326 | 16788 | 15090 |
| **Number of points in DVC point cloud** | 1363157 | 1069502 | 702778 |

**Supplementary Table 2 –** Parameters and their default values implemented in *TomoSAXS* reconstruction.

| **Parameter** | **Default value** | **Units** | **Notes** |
|---|---|---|---|
| minimum χ | 0 | degrees | Minimum χ for sampling/simulating scattering |
| maximum χ | 180 | degrees | Maximum χ for sampling/simulating scattering |
| nchi | 180 | n.a. | Binning number for χ sampling |
| q0_m | 0.28 | $nm^{-1}$ | Median $q_0$ value for simulation |
| q0m_low | 0.262 | $nm^{-1}$ | Minimum $q_0$ value for simulation |
| q0m_high | 0.302 | $nm^{-1}$ | Maximum $q_0$ value for simulation |
| nq0 | 50 | n.a. | Binning number for $q$ sampling |
| threshold_interference | 10 | % | Threshold maximum percentage value of total combined scattering that neighbouring fibres must contribute for single fibre scattering under-analysis to be considered independent. |
| threshold_detection | 3.50E+06 | Intensity | Minimum absolute level of scattering intensity threshold for considering single/combined scattering. |
| chi_range_sample | 3 | degrees | Radius across χ for independent scattering subsamples |
| nslices | 3 | n.a. | Number of independent scattering subsamples |
| wavelen | 0.08856 | nm | Experimental wavelength |

| | | | |
|---|---|---|---|
| q2i | 0.262 | $nm^{-1}$ | Minimum $q_0$ value for inner background correction of radial integrations |
| q1i | 0.267 | $nm^{-1}$ | Maximum $q_0$ value for inner background correction of radial integrations |
| q1o | 0.302 | $nm^{-1}$ | Minimum $q_0$ value for outer background correction of radial integrations |
| q2o | 0.312 | $nm^{-1}$ | Maximum $q_0$ value for outer background correction of radial integrations |
| bg_start | 0.2 | $nm^{-1}$ | Minimum $q_0$ value for sampling/simulating azimuthal integrations |
| bg_end | 0.35 | $nm^{-1}$ | Maximum $q_0$ value for sampling/simulating azimuthal integrations |
| q0_sim | 0.28 | $nm^{-1}$ | Initial $q_0$ estimate for 3D diffraction model fitting |
| wa_sim | 0.0056 | $nm^{-1}$ | Initial $w_a$ estimate for 3D diffraction model fitting |
| wMu_sim | 0.2 | $nm^{-1}$ | Initial $w_\mu$ estimate for 3D diffraction model fitting |
| delta_sim | 5.00E-01 | n.a. | Initial $\delta$ value for 3D diffraction model fitting |
| q0_min | 0.255 | $nm^{-1}$ | Minimum $q_0$ estimate for 3D diffraction model fitting |
| q0_max | 0.314 | $nm^{-1}$ | Maximum $q_0$ estimate for 3D diffraction model fitting |
| wa_min | 0.0005 | $nm^{-1}$ | Minimum $w_a$ estimate for 3D diffraction model fitting |
| wa_max | 0.01 | $nm^{-1}$ | Maximum $w_a$ estimate for 3D diffraction model fitting |
| wMu_min | 0.02 | $nm^{-1}$ | Minimum $w_\mu$ estimate for 3D diffraction model fitting |
| wMu_max | 0.8 | $nm^{-1}$ | Maximum $w_\mu$ estimate for 3D diffraction model fitting |
| amp_min | 1.00E+03 | intensity | Minimum amplitude estimate for 3D diffraction model fitting |
| amp_max | 6.00E+09 | intensity | Maximum amplitude estimate for 3D diffraction model fitting |
| min_error | 25 | % | Threshold minimum value for normalised standard error for fit quality |
| min_fit_q0 | 0.256 | $nm^{-1}$ | Threshold minimum value for estimated $q_0$ |
| max_fit_q0 | 0.313 | $nm^{-1}$ | Threshold maximum value for estimated $q_0$ |
| min_fit_wa | 0.0006 | $nm^{-1}$ | Threshold minimum value for estimated $w_a$ |
| max_fit_wa | 0.0098 | $nm^{-1}$ | Threshold maximum value for estimated $w_a$ |
| min_fit_wMu | 0.06 | $nm^{-1}$ | Threshold minimum value for estimated $w_\mu$ |
| max_fit_wMu | 0.795 | $nm^{-1}$ | Threshold maximum value for estimated $w_\mu$ |
| diam_min | 5 | nm | Threshold minimum value for estimated fibril diameter |
| diam_max | 200 | nm | Threshold maximum value for estimated fibril diameter |
| searchwindow | 10 | degrees | Search window across $\chi$ for isolating combined scattering fibres |
| thresh_detect | 3.50E+06 | intensity | Minimum absolute level of scattering intensity threshold for considering combined (2-3 fibres) independent scattering. |
| thresh_combined | 80 | % | Minimum proportion of total combined scattering for combined (2-3 fibres) scattering to be considered independent |
| fitwindow | 10 | degrees | Minimum distance across $\chi$ for combined scattering fibres to provide scattering above minimum % threshold |
| thresh_individual | 20 | % | Minimum percentage of individual fibres to provide when combining scattering of multiple (2-3) fibres |

| | | | |
|---|---|---|---|
| Noise_fac | 1 | n.a. | Noise modifier (simulated background scattering divided by this value) |
| dq0_m | 0.022 | n.a. | Bounds for $q_0$ randomizer for digital phantom simulations |
| dwa_m | 0.003 | n.a. | Bounds for $w_a$ randomizer for digital phantom simulations |
| dwMu_m | 0.9 | n.a. | Bounds for $w_\mu$ randomizer for digital phantom simulations |
| amp_m | 5000 | n.a. | Bounds for amplitude randomizer for digital phantom simulations |

**Supplementary Figure 1 – Sample environment.** Diagram of the adapted Deben CT 500 mechanical testing rig with 3D printed sample holder and phosphate-buffered saline (PBS) fluid delivery tube.

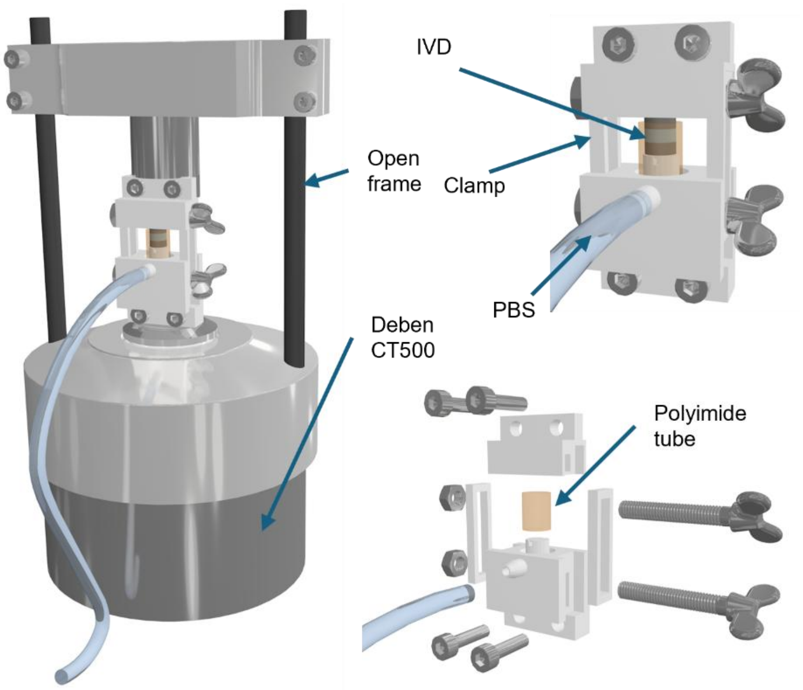

**Supplementary Note 1 – Radiation dose calculation for tomographic SAXS**

Per-beam-path radiation dose for SAXS scans was calculated using the method described in Badar et al. *Adv. Science* (2025)[1], modified here to account for tomographic scanning:

$$Dose\ (Gy) = \left( \frac{Incident\ flux(ph/s) * exposure\ time\ (s) * absorption\ ratio * X - ray\ energy\ (j)}{sample\ volume\ along\ beam\ (m^3) * sample\ density\ (\frac{kg}{m^3})} \right) * no.rotations \tag{1}$$

Incident flux was here $3.03 \times 10^{10}$ *ph/s*, calculated as the proportion of original flux ($3.70 \times 10^{10}$ *ph/s*) not absorbed by PBS solution surrounding the sample. This was calculated using the linear X-ray attenuation coefficient (2.00 cm), and average distance travelled through PBS before the sample (1 mm) (X-ray attenuation = $e^{(2.00*(1*0.1))}$ = 0.819) (non-attenuated flux = $3.70 \times 10^{10}$ * 0.819 *ph/s*). Estimated X-ray absorption ratio was 0.3 and X-ray energy was $3.24 \times 10^{-15}$ *J*. Average sample volume along the beam was $3.42 \times 10^{-12}$ $m^3$, and sample density was estimated as 1100 kg/$m^3$. This provided an estimated dose per-beam-path of 0.81 kGy. With nine rotations applied, this resulted in a total per-beam-path dose of 7.3 kGy per-scan.

**Supplementary Note 2 – Radiation dose calculation for pCT**

Radiation dose for each high-resolution scan was calculated using previously described methods[2,3]. Dose was estimated as

$$Dose\ (Gy) = \frac{Exposure\ time\ (s) \times Absorbed\ energy\ flux\ (\frac{J}{sm^2})}{Mass\ per\ unit\ area\ (\frac{kg}{m^2})} \tag{2}$$

Total exposure time is given by the number of projections multiplied by 0.15 s exposure time (1801*0.15 = 270.15 s). Using an absorbed energy flux of $4.9 \times 10^{-4}$, mass per unit area of 1 $gcm^3$, and thickness of 5 mm, total dose for one scan was calculated as 26.6 kGy.

The 7 mm ID gap used for low-resolution scans resulted in a 95 % reduction in flux compared to high-resolution scans, giving a total dose of 1.33 kGy.

**Supplementary Note 3 – Fibre tracing**

Fibres were segmented from the pCT images using the XFiber module in Avizo 3D (v2023.2). Cylinder Correlation was applied with a cylinder length of 40 voxels (65 µm), angular sampling 5, mask radius 3 voxels (4.9 µm), outer radius 2 voxels (3.2 µm), inner radius 0, and bright-on-dark contrast. Correlation and orientation fields were masked prior to tracing to minimise artefactual fibres. The annulus fibrosus was manually segmented every 50th slice with interpolation, and interlamellar space excluded by interactive thresholding. Trace Correlation Lines was then applied with the following parameters: minimum seed correlation 80, minimum continuation quality 60, direction coefficient 0.2, minimum fibre spacing 5 voxels, search cone length 40 voxels, search cone angle 37°, and minimum step size 10%. The minimum fibre length threshold was sample-specific, set according to IVD height (Supplementary Table 1), to balance exclusion of artefacts with retention of true fibres.

**Supplementary Note 4 – Fibre orientation calculations**

Fibre orientation was calculated using custom-written Matlab scripts. The fibre identification number was defined as the Segment ID number output from the Avizo spatial graph and was conserved throughout all analyses. Point coordinates output from Avizo were used to identify fibre direction, then points were placed every 4 voxels (6.5 μm) along each fibre. The coordinates of these points were saved as a text file for input into DVC analysis. A $3^{rd}$ order polynomial curve was fit to the points for each fibre, and the tangent to the curve calculated at each point. The angle between the tangent and the vertical (α) and horizontal (β) axes were calculated between $0^\circ$ and $180^\circ$. Curve fitting parameters were saved as a data matrix for further analysis. The x, y, z coordinates, fibre ID, α and β orientations for each point were saved as a .csv file for further processing in Avizo.

**Supplementary Note 5 – Fibre ID and orientation image volume generation**

The Avizo spatial graph file output from fibre tracing was converted to a binary label image with a voxel size of 8.125 μm (5 pCT voxels). Fibre ID, α, and β values were converted to 3D volume images with the same dimensions as the fibre label image, using the nearest-neighbours method for interpolation. Fibre ID was converted to a 16-bit image and α and β were converted to 8-bit images using a 1-to-1 scaling. These images were then masked with the fibre label and exported as 3D tiffs for input into the *TomoSAXS* reconstruction.

**Supplementary Note 6 – Radial and circumferential strain calculation**

Tissue-level radial and circumferential strains were calculated using the Python (3.12) script, which projects the Lagrangian strain tensor at each point along the local radial and circumferential direction vectors. A smooth manual segmentation of the AF was performed in the xy plane every 100 slices in z and linearly interpolated in Avizo to produce a 3D binary mask, which was saved as a 3D tif file. This binary mask, along with the tissue-level strain data file output from DVC, were imported into Python. Inner and outer contour lines were fit to the AF mask in each xy slice, a midline was placed between the contour lines and two further lines added between the midline and inner contour, and midline and outer contour (Extended Data Fig. 3a). Radial and circumferential vectors were calculated for 1000 points along each contour line. For every point in the DVC point cloud, the vectors at the closest point on a contour line were used for strain projection, and the Lagrangian strain tensor projected along these vectors.

**Supplementary Note 7 – Fibre strain calculation**

A Matlab script was used for fibre strain calculation. Fibre strain was calculated from a variable degree polynomial fit to displacements of points along their associated fibre direction. Polynomial degree varied according to fibre length, with the lowest degree set to 2 (quadratic) and an extra degree added per 320 voxel (520 μm) fibre length up to a maximum polynomial degree of 9. This was done to reflect the increasing complexity of strain fields felt by longer fibres (Supplementary Figure 2).

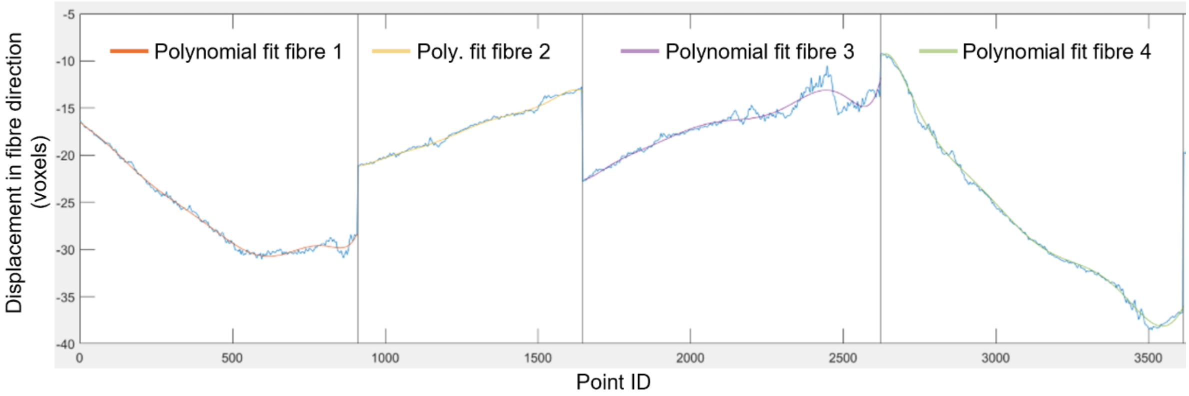


**Supplementary Figure 2 – Polynomial fitting for fibre strain calculation.**

**Supplementary Note 8 – Distance between endplates and bulk IVD strain calculation**

Segmentation and digital volume correlation (DVC) of the vertebral endplates was performed according to methods described in Parmenter et al 2025[3]. Mean distance between the endplates, listed in Table S1, was calculated as the distance between the mean x, y, z coordinates of DVC point clouds created from the two endplates before loading. Distance after loading was calculated by applying the DVC measured u, v, and w displacements to the endplate point clouds and calculating the new mean distance. The change in mean distance before and after loading was used to calculate bulk compressive strain applied to the IVD. The minimum distance between the endplates was calculated using a custom-written Matlab script (Matlab R2023a) which uses the *knnsearch* function to find the distances between all points in the cranial and caudal endplate point clouds and then calculate the minimum distance between two points. The new distance between these points after loading was calculated by applying the DVC displacements to the points. The percentage change in distance at the minimum distance between the endplates was used to approximate the maximum strain applied to the IVD.

**Supplementary Note 9 – DVC accuracy calculation**

The double projection zero strain method[2,3] was used to quantify the accuracy of DVC displacements and strains. A 'sacrificial' whole IVD sample was used for double (3600) projection scanning; odd and even projections were reconstructed separately, resulting in two 1800 projection scans taken at the same time, removing sample viscoelastic effects from DVC error calculations. Fibre tracing and DVC point cloud generation for the entire annulus fibrosus was performed as described above, along with DVC and strain calculations.

Displacement and strain accuracy and precision were calculated using the mean absolute error (MAER) and standard deviation of error (SDER)[3,4]:

$$MAER = \frac{1}{N}\sum_{k=1}^{N}(\frac{1}{6}\sum_{c=1}^{6}\left|\varepsilon_{c,k}\right|) \quad \textbf{(3)}$$

$$SDER = \sqrt{\frac{1}{N}\sum_{k=1}^{N}(\frac{1}{6}\sum_{c=1}^{6}\left|\varepsilon_{c,k}\right| - MAER)^{2}} \quad \textbf{(4)}$$

Where N represents the number of points, k is the point in the point cloud, c is the component of strain, and ε is the strain value (analogous calculations performed for components of displacement).

Overall zero-strain displacement accuracy was 99.4 nm and precision 148 nm. Tissue strain accuracy was 0.166 % and precision 0.234 %. Fibre direction displacement accuracy was 112 nm and precision 181 nm, this is larger than the overall displacement accuracy due to the ‘long valley’ optimisation problem associated with tracking fibres, where there are fewer unique landmarks along the length of a fibre, so it is easy for points to slide along the fibre direction during the optimisation process. Fibre strain accuracy was 0.07 % and precision 0.15 %.

The ability of the DVC analysis to track deformation was further tested by applying a virtual compression of 10 % in the z direction to the second image generated from the double projection method. DVC measurements of displacement and strain were then compared to the actual virtual deformation applied to the image, to again determine accuracy (MAER) and precision (SDER). Overall 10 % strain displacement accuracy was 198 nm, precision 1.41 µm. Tissue strain accuracy was 0.399 % and precision 2.77 %. Fibre direction displacement accuracy was 219 nm, precision 1.72 µm. Fibre strain accuracy was 0.453 %, precision 2.16 %.

**Supplementary Note 10 – Loaded fibre orientation calculation**

To calculate orientation for the same fibres in each sample after loading, DVC displacements were applied to the fibre points in Matlab. Orientation at each point in the fibre was calculated the same way as for preloaded images using a $3^{rd}$ order polynomial fit. The fibre mask used for creating loaded Fibre ID, α, and β TIFFs was generated by creating a point cloud with points placed every voxel along the loaded fibres and converting this geometry to a label image in Avizo. The updated orientation TIFF stacks were then used for loaded data *TomoSAXS* reconstructions.

**Supplementary Note 11 – 3D diffraction Modelling**

The SAXS signal for the meridional reflections in 3D was modelled using a similar approach to our recent publication[1]. In this work, each reflection was modelled as a 3D highly oblate ellipsoid in reciprocal space. The ellipsoid had its thin axis along the fibre direction with radial width $w_a$ and a much larger lateral width $w_p$, where fibril diameter and $w_p$ have an inverse relationship which can be calculated numerically (Figure S8 in Supplementary Information of W. Badar et al[1]). In our scattering data from IVD collagen in the current work, we observed that the ellipsoids were not perfectly flat but not curved enough to be spherical sectors. To reflect this, each reflection was modelled as a partly curved ellipsoid characterised by a δ-parameter (δ=0: flat; δ=1: spherical sector). The fibril diameter was obtained by applying the inverse relation between $w_p$ and fibril diameter, where $w_p$ was calculated as $q_0 \tan(w_\mu)$ with $w_\mu$ a fit parameter in the model.

**Supplementary Note 12 – Pre-processing of pCT data for *TomoSAXS* analysis**

Several pre-processing steps are required before the operation of the *TomoSAXS* reconstruction pipeline. First, as the fibre tracing procedure involves subsampling (only considering a portion of information-rich pCT data) and down-sampling (resultant fibre tracing datasets have a larger voxel size than original pCT datasets), fibre tracing datasets must be padded to return them to the same absolute volume as the original pCT data. Second, due to the aspect ratio of our samples and the necessity of including the surrounding kapton tube for spatial registration (see below), our high resolution pCT data (1.625 μm voxel size) has to be spatially calibrated with data from a lower resolution (2.6 μm voxel size) scan of each sample.

This process follows:

*1. Estimation of padding volumes for fibre tracing data.* The necessary information for padding fibre tracing data is provided as an output from the Avizo software. This comprises information on the number of voxels in the outputted fibre tracing dataset, the physical size of this dataset (in terms of original pCT voxels), and its location with respect to the original pCT volume. This data is used to calculate the amount of padding (empty voxels) needed to apply to the fibre tracing data for it to replicate the original aspect ratio of the pCT volume. Values of padding proportions to be applied before/after the fibre tracing dataset across each axis are saved in a .csv file. Padding is later applied together with the spatial calibration of the original high and low resolution pCT data (see below).

*2. Scaling CT data.* The high resolution and low resolution CT reconstructions are spatially down-sampled to 6.5 μm voxel size in ImageJ/Fiji[5] (image>adjust>size), and the resultant datasets saved as 8-bit tiff stacks. A shared region found in a single slice is then isolated in both down-sampled datasets and segmented, saving the segmented slice with its slice number as the file name. The difference in absolute slice position of the segmentation is also saved (i.e. the region will be in a higher slice number in the down-sampled lower resolution reconstruction).

*3. Padding and spatial calibration.* This is accomplished algorithmically using the "ct_reg.py" script. This uses the padding values saved for fibre tracing data (step 1) to create empty arrays along each axis, concatenating them to the fibre tracing volume. The centre of the regions segmented from both low and high resolution pCT reconstructions are then found, and used to overlay the high resolution data onto the low resolution data, based on the offset found between the slice numbers in the regions respective positions in each volume. This spatially calibrated volume is saved as a new dataset in both 16 bit and 8 bit.

*4. Creation of pCT map for comparison with mapping WAXS map.* A further pre-processing step involves creating a 2D map of the sample in pCT data that aligns with the mapping scan taken before SAXS scanning (Extended Data Fig. 1). The calibrated pCT volume created by "ct_reg.py" is opened in Fiji, and resliced longitudinally (image>stacks>reslice), creating a new stack orientated along the 0° of the pCT scan (the same orientation as the map). This is then z-projected (image>stacks>z-projection), to create a single image of averaged values through the longitudinal volume, to compare spatially with WAXS maps as part of the 3D registration process (see below).

*5. Processing Kapton data.* To use the Kapton tube in the spatial registration process, it must first be isolated in pCT datasets. This is performed using the "ROI manager" in Fiji, by segmenting the tube every 50 slices in the calibrated pCT dataset, and interpolating between them. This creates a segmented dataset of the same aspect ratio as the original data, comprising only information for the Kapton tube.

*6. Segmenting samples in pCT data.* A final pre-processing step is performed here, to provide variable sample thickness estimates for background correction of SAXS data. Padded fibre tracing data is incorporated into calibrated pCT data in Fiji by summing the greyscale values of each dataset, creating volumes where both bone and soft tissues are represented by greyscale values significantly above those of the background. This allows automatic thresholding of sample versus background in Fiji, and thresholded values are saved as binary masks (removing the Kapton data through multiplication with inverted kapton segmentation datasets).

**Supplementary Note 13 – Spatial co-registration of pCT and SAXS data**

*TomoSAXS* scans are acquired by rotating a sample over a set number of orientations over a complete 180° rotation. Each orientation is represented by a two dimensional raster map of SAXS frames, that includes the edges of the Kapton tube around the sample. The centre of this rotation is independent of the rotation of the corresponding CT scan from-which fibre orientations are estimated. These datasets must be registered both vertically and horizontally, for each raster map representing each orientation of the *TomoSAXS*TomoSAXS scan, in order for the identity of isolated fibres in CT data to be consistently retained through the *TomoSAXS*TomoSAXS scan.

First, perquisite data is loaded by the user using a series of graphical user interfaces (GUIs). This loads:

1. *"Scan name"* - the name given to the *TomoSAXS*TomoSAXS scan in the accompanying fibre tracing "vix_padding" excel file.
2. *"Original CT data"* - the folder containing the original CT data.
3. *"Inverted resliced CT map"* - the file comprising the resliced, grayscale inverted CT map corresponding to the WAXS map used for registration.
4. *"Kapton CT dataset"* - the folder containing the segmented Kapton tube data.
5. *"Beta/phi fibre tracing data"* - the folder containing the (original unpadded) beta/phi fibre tracing data.
6. *"Alpha/theta fibre tracing data"* - the folder containing the (original unpadded) alpha/theta fibre tracing data.
7. *"WAXS map data"* - the .nxs file of the WAXS map.
8. *"Output folder"* - the folder that the user wishes to output data generated by the registration script (example figures and tables).
9. *"Original CT voxel size (um)"* - The voxel size of the original CT data in microns.
10. *"Inverted CT voxel size (um)"* - The voxel size of the inverted CT data (may be adjusted if processed on a laptop due to limited computing power).
11. *"Kapton data voxel size (um)"* - The voxel size of the Kapton segmented data (may be adjusted if processed on a laptop due to limited computing power).
12. *"Fibre tracing voxel scale"* - The downsampling scale used for fibre tracing data creation.
13. *"Kapton tube diameter (um)”* - diameter of kapton tube in microns.

14. *"SAXS rotational direction"* - direction of rotation for TomoSAXS scan.
15. Original SAXS datasets (.nxs fles)
16. *"Number of rotational angles in TomoSAXS scan".*
17. *"start angle"* - axis orientation of the first orientation of the TomoSAXS scan.
18. *"end angle"* - axis orientation of the last orientation of the TomoSAXS scan.
19. *"angle of WAXS map"* – axis orientation that the coarse mapping scan (Extended Data Fig. 1) was conducted at.

Once these inputs have been finalised, the mapping WAXS (wide angle X-ray scattering – conducted in parallel to SAXS acquisition) map (Extended Data Fig. 1) is generated, and the user must specify a sample-specific morphological landmark. The lowest vertical point of the upper vertebral endplate is usually specified here (for rodent IVD samples). Selection of this landmark generates the z-projected pCT map (see Pre-processing of pCT data for SAXS analysis), where the same region is selected. This allows estimation of the absolute height within the pCT dataset that is represented by the *TomoSAXS* scan, and isolates the pCT, kapton segmentation, sample thickness data, fibre tracing, and DVC data for this region.

Next, the fibre tracing data represented by the *TomoSAXS* slices is further pre-processed and padded to overlap with the SAXS data in each orientation of the SAXS tomography. This is performed independently for each rotation (as raster maps were collected independently during the SAXS scan). Calibration is performed by rotating the kapton tube data by the respective offset from 0° for each scanning orientation, and calculating the resultant position of the left-hand edge of the tube along the x-axis. This edge is also found in the respective SAXS data, by identifying peaks in total SAXS scanning across the scan - provided by characteristically intense scatter experienced at the edge of the kapton tube. Identification of this edge in both datasets allows fibre tracing data (and sample thickness data), rotated alongside the Kapton segmentation, to be further padded along it’s x-axis until the edge is found at the same point along this axis in both datasets.

The registered, padded fibre tracing data is subsampled across the X-axis in accordance with the difference in absolute voxel size between fibre tracing (8.125 µm) and SAXS data (20 µm). For each SAXS beam-path, the respective fibre tracing data for each fibre encountered along the beam-path are isolated and added to a dataset (index, vertical angle, horizontal angle). Rotated sample thickness data is also subsampled, with the distance between the first and last sample voxel across the beam-path used to estimate the thickness of the encountered sample. Finally, as SAXS X-ray beams are not perfectly collimated and scattering may not be restricted to fibres within the confines of sub-sampled beam-paths, a weighting factor is applied to all fibres within the immediately surrounding beam-paths based on proximity to the beam centre:

$$weighting\ factor = e^{-d^2/2w^2} \quad \textbf{(5)}$$

where d is the distance from beam centre and w the beam diameter.

**Supplementary Note 14 – Estimation of scattering amplitude using singular value decomposition**

Before spatially registered datasets can be implemented for *TomoSAXS* reconstruction, an amplification factor must be applied between modelled scatter and measured scatter for individual fibres. 3D diffraction models produce estimates of relative scattering intensity for fibres based on their angular orientation and applied scattering parameters. These values are several orders of magnitude lower than measured scatter in our SAXS data. To overcome this, and provide more realistic initial estimates of amplitude for improving fits between modelled and measured scatter for real individual fibres, we estimate an amplitude factor for models by applying singular value decomposition (SVD) between simulated and observed scattering data measured across the circumferential χ axis.

SVD (alongside *TomoSAXS* fitting) is partitioned into single horizontal *TomoSAXS* slices to maximise computational efficiency by running the "svd_module.py" program (GITHUB LINK) independently for each slice using cluster computing. For each slice, observed scattering data for each SAXS beam-path is interpolated across the χ axis (-90° – 90° to minimise the effects of masking over the 3rd order region of *q*). Interpolations are conducted across the scattering region (*q*: 0.26-32 $nm^{-1}$), and immediately inside (*q*: 0.22:0.26 $nm^{-1}$) and outside (*q*: 0.32:0.36 $nm^{-1}$) of this region, with mean values created between these data to correct for background scatter. Background corrected I(χ) integrations are further smoothed using a Savitzky-Golay filter (filter window = 9; polynomial order = 1). These are concatenated into a single array for statistical comparison with modelled data.

Model data is created for each beam-path by isolating their composite fibres and simulating per-fibre scatter across the same χ region as observed data using their orientation (relative to the orientation of the X-ray beam) and weighting data. Scattering parameters are provided using mean values estimated from measurements of assumed independent fibre scattering of the measured data (Supplementary Table 2). Simulations are concatenated into a two-dimensional array, with rows defined by fibre index and columns defined by the same point along I(χ) integrations across beam-paths (identical to those of the measured beam-paths).

The measured and simulated arrays are finally statistically correlated using non-negative linear regression analysis, providing initial estimates of amplification factors for individual fibres. The majority of fibres (~80%) provide non-negative estimates of scattering amplitude. However, for others with weaker observed scattering, issues arise from negative values in background-corrected observed I(χ) data and the use of non-negative linear regression during SVD – resulting in 0 values for estimated amplitude. All values are saved, alongside the index, orientation, and weighting data for each fibre (respective to the beam-path orientation), for sub-entries specific to each beam-path within entries for each orientational angle of the SAXS tomography in a Python dictionary. This dictionary is used in *TomoSAXS* fitting.

**Supplementary Note 15 – *TomoSAXS* fitting**

For each horizontal *TomoSAXS* slice, SVD analysis provides Python dictionaries partitioned into each orientational angle of the SAXS tomography. These entries are composed of sub-entries for each X-ray beam-path that consist of the orientation, index, weighting, and initial estimate of scattering amplitude for sampled fibres. This allows for simulation of the SAXS tomography and isolation and fitting of the independent scattering of individual fibres within single beam-paths.

Fitting is performed using a central recon_library.py Python library. This simulates scattering across χ from each unfitted fibre in a single beam-path using their 3D orientation, weight, initial estimated scattering amplitude, and mean scattering parameters (Supplementary Table 2) (fitted fibres are not considered here as their contribution is accounted for during fitting – see below). Partitioning of these fibres allows isolation of the individual contribution of the fibre to combined scatter and testing whether this contribution exceeds a user-defined proportion of total scatter (default = >90%) for a user-defined minimum portion across χ (default = >10°). The rotation, beam-path index, fibre index, and region of independent scatter are added to a list, outputted as a return of the primary library function.

Dependent on user-defined inputs (bullion True/False for solving), this region of χ is sampled in the observed SAXS frame for the beam-path, following a series of tests. These first consider the portion of independent scattering space that is masked (represented by empty pixels between sub arrays of the I22 Pilatus detector). Only un-masked positions are considered for analysis. If a user-defined minimum (default = >10°) remains un-masked, and this portion comprises scatter from solved fibres, the combined scatter from solved fibres (simulated using their fitted parameters) across this region is compared to the simulated scatter for the fibre under analysis. Only fibres that provide a proportion of scattering above a user-defined threshold (default = 40%) compared to combined solved fibre scattering are considered for further analysis. These fibres are finally sampled across *q* within equally spaced and equally sized sub-regions within the remaining unmasked independent portion of χ (default = 6°).

Sampled I($q$) integrations are fitted with 3D scattering models to provide estimates of their scattering parameters. These models are informed by parameter objects consisting of scattering parameters for the fibre under-analysis and any fibres previously fitted within the beam-path. Values for the fibre under-analysis are providing from mean values used for independence estimation, that are allowed to be optimised within certain bounds (Supplementary Table 2), alongside their orientation, weight, and index data (not allowed to change during model fitting). Previously fitted fibres are provided with their estimates for each scattering parameter, which are not allowed to change during model fitting. Models thus consider combined simulated scattering for each fibre in the parameter object. Parameters for the fibre under-analysis are optimised using the Nelder-Mead method to provide best-fitting values. These are further tested to check the quality of fits. First, a series of threshold values for each scattering parameter are applied, failure to meet one-of-which results in rejection of the fit (Supplementary Table 2). For fits that pass these thresholds, their standard error (SE) is estimated by using the fitted parameters (for each fibre in the parameters object) to model scattering for each individual I(q) sample and comparing simulated to observed values (normalised to the highest intensity value in both simulated and observed data). If one sample provides an SE value below a user-defined threshold (default = 25), the fit is deemed as successful and the parameter values are added to each instance of the fibre under-analysis in the dictionary.

Nanoscale parameter values (mean fibril D-period, D-period variation, fibril diameter) are estimated from fitted scattering properties. Mean D-period is calculated from the estimated fundamental scattering vector ($q_0$), with respect to the harmonic order under study (default 3rd order approximately 0.26-0.31 $nm^{-1}$):

$$D = (2 * \pi) * 3/q_0 \quad \textbf{(6)}$$

Where D equals mean fibril D-period (nm).

D-period variation (D variation; nm) is calculated from the estimated width of the peak along $q$ ($w_a$), following:

$$min.D = (2 * \pi) * 3/(q_0 + w_a) \quad \textbf{(7)}$$

$$max.D = (2 * \pi) * 3/(q_0 - w_a) \quad \textbf{(8)}$$

$$D\ variation = max.D - min.D \quad \textbf{(9)}$$

Fibril diameter is estimated using the width of the peak along $\chi$ ($w_p$), which is itself calculated from the estimated Cartesian width of the peak ($w_\mu$) and $q_0$:

$$w_p = q_0 * \tan(w_\mu) \quad \textbf{(8)}$$

Fibril diameter (nm) is then estimated from $w_p$ using the regression formula between these properties between measured diameter and $w_p$ by W. Badar et al, *Adv. Science* (2025):

$$fibril\ diameter = \left(2.857 * \left(\frac{1}{w_p}\right)\right) + 10 \quad \textbf{(9)}$$

If a single fibre is not found to provide sufficient portions of independent scatter, it can be related to neighbouring fibres to test whether two, or a (current) maximum of three fibres can be fitted simultaneously. Dependent on user-defined inputs (bullion True/False for combined solving), the simulated scattering of the fibre under-analysis is combined with scattering of each unsolved fibre in the respective beam-path and their combined scatter compared to total scattering. If their combined scattering meets the requirements for definition as independent (Supplementary Table 2), their region of shared independence is sampled using $I(q)$ integration and fitted by adding entries for each fibre in the parameters object. An additional stipulation is added for three fibres, via additional analysis of the proportion of estimated scatter from each fibre. Only cases where each fibre represents scattering intensity above a user-defined proportion of combined scattering (default = 20%) are considered for fitting. This is to avoid potential overprinting of low intensity fibres from neighbouring fibres during fitting.

**Supplementary Note 16 – *TomoSAXS* Fitting procedure**

The above method is applied sequentially under differing conditions through the *TomoSAXS* reconstruction process. In the first iteration, only scattering instances where single fibres with an amplification factor >0 are estimated to provide independent scattering are considered. This is performed within a "while" loop that performs single assessments of the complete SAXS tomography without inline fitting and no analysis of combined scatter. From this, output lists of potential independent scattering events for single fibres from each iteration are obtained, which are subsequently fitted sequentially (with successful fits of single fibres allowed to contributed to analyses of subsequent fibres). For fibres where multiple independent instances are estimated from one assessment of the SAXS, tomography, each instance is sequentially investigated in descending order of estimated scattering intensity. This loop is closed once every incidence of independent scatter has been investigated.

Next, fitting is permitted for single, double and triple cases in-line with the SAXS tomography simulation. The simulation is repeated to a user-defined limit (default = 10), or until no new fits are provided during an entire simulation.

Finally, fibres that provided an amplification factor of 0 (see Estimation of scattering amplitude using singular value decomposition) are provided with an amplification factor based on the mean fitted amplitude found for all fitted fibres. The simulation program with in-line fitting of single, double, and triple cases is then repeated. This provides a maximum possible number of fits for individual fibres in the sample, based on the fibre geometry and experimental parameters.

**Supplementary Note 17 – Per-fibre nanoscale strain calculation**

DVC vertical (z-axis) displacement values were used to located the position of fibres in the loaded TomoSAXS reconstruction. 3D image volumes of z-displacement were generated the same as for orientation (see Supplementary Note 5). These maps provide estimates of vertical deformation in each voxel, calculated at the same resolution as the original pCT data (1.625 µm voxel size); while the maps used for *TomoSAXS* reconstruction are based on fibre tracing data (down-sampled to 8.125 µm voxel size). The potential inaccuracy introduced by this down-sampling was further enhanced in cases where the estimated position of the fibre point in the loaded *TomoSAXS* scan was at the edge of two 20 µm *TomoSAXS* slices. To account for this, a search window was established to find the matching loaded position for each unloaded point, with the minimum and maximum *z* axis positions set to the estimated deformation point minus or plus four *s*CT voxels respectively. If this window crossed the boundary between two *TomoSAXS* slices, and a fit was provided for the fibre in both slices, the slice with the closest estimate of D-period to the unloaded value was considered as the matching loaded point.

**Supplementary Note 18 – Digital phantom simulation**

Randomised fitted values for nanoscale properties from *TomoSAXS* fitting (Extended Data Fig. 5) are used to provide artificial scattering parameters for each fibre. Parameter estimates from fitted fibres are randomly assigned to all fibres through random selection of their index. These values are further changed through the application of random noise to these values (Supplementary Table 2). For each fibre, artificial values are assigned to all instances in the *TomoSAXS* fitting dictionary, preserving their orientation, weighting and initial amplitude estimate.

Artificial parameters are processed with the same Python library as used in *TomoSAXS* fitting of real scattering data. A major modification for analysing artificial data is performed in the sampling function for real data (Extended Data Fig. 7a). For artificial data, tests for masked versus unmasked scattering regions are based on the real SAXS frame representative of the fibre's beam-path (Extended Data Fig. 7ai). As for sampling real data, simulation testing for overprinting from fitted fibres applies their fitted values, and mean values for unfitted fibres under analysis. However, if fibres pass both of these tests then "sampling" of their scattering is created using I($q$) integration of 3D diffraction models across independent scattering sub-regions (Extended Data Fig. 7aii,aiv), with intensity values dictated by their artificial scattering parameters. To replicate noise found in real scattering data, the SAXS frame representative of the beam-path is also sampled across the same sub-regions, and background scattering data collected across $q$ inside of the 3$^{rd}$ order region of collagen scattering ($q$ = 0.20-0.25 nm$^{-}$

$^{1}$) (Extended Data Fig. 7aiii). This data is divided by a modification parameter (default = 1) to modulate the proportion of noise implemented to simulated data (Extended Data Fig. 7av). The proportion of the modulated background data's standard deviation to the mean intensity value of the complete sample is calculated. This value is multiplied by an array of random float values between -1 and 1 to create a noise function that is added to the original simulation for each independent sub-region (Extended Data Fig. 7avi).

Noisy simulated data is subsequently fitted using the same procedure as for real data. Implementation of this fitting procedure follows Supplementary Note 14. We find strong correlation between known and estimated values for digital phantoms of each IVD sample ($r^2$ = 0.641-0.894; $p$<0.001) (Extended Data Fig. 7b). These provide maps of each property that capture randomised patterns in known properties. Regression slopes between known and estimated properties are below 1:1, suggesting that lower known values are slightly over-estimated and higher values underestimated using *TomoSAXS.* The underestimation of higher values is most pronounced for fibril diameter (Extended Data Fig. 7diii). However, while absolute spatial contrast is lower in our estimated maps of nanoscale properties, trends in relative values are maintained (Extended Data Fig. 7b)

**Supplementary Note 19 – Comparison between two-dimensional SAXS and *TomoSAXS* analysis of IVD**

To understand the potential advantages of the *TomoSAXS* technique compared to traditional two-dimensional (2D) SAXS analysis of soft collagenous tissues, we also applied 2D SAXS analysis to SAXS raster maps conducted at the 0° rotation for each of our IVD samples.

Analysis was performed by fitting Gaussian models to integrated intensity ($I(q)$) profiles across $q$ between 0.2-0.35 nm$^{-1}$ and across $\chi$ between 0-180°. D-period (nm) was estimated per beam-path from the estimated centre-point ($q_0$) of resultant models along $q$ (Equation 6) (Extended Data Fig. ai) and d-period variation derived from their σ ($w_a$) (Equations 7-9) (Extended Data Fig. aii). As no single fibre deconvolution was applied here, values for D-period and D-period variation can only be considered as average values for all fibres within the respective beam-path.

This results in significant differences between D-period values estimated using 2D SAXS compared to *TomoSAXS* for each sample (Extended Data Fig. 6ai,iii). Measurements of convoluted scattering across beam-path frames in 2D SAXS result in variation (standard deviation) in estimated D-periods that is considerably lower than for *TomoSAXS* (Kruskal-Wallis $H$ = 8.208, $p$ = 0.004) (Extended Data Figs 6ai,iii). Mean per-sample D-period values are also lower than for *TomoSAXS* (Kruskal-Wallis $H$ = 8.308, $p$ = 0.004). This is likely due to the inability of 2D SAXS to account for potential orientation effects of fibres whose meridional scattering peaks divert from ellipsoidal in shape (Δ<0.99). As these fibres prograde away from vertical alignment (α = 0), their orientation effects their intersection with the Ewald Sphere, which is observed as an increase in the effective position of scattering along $q$. Measurements of observed scatter across $q$ may thus provide $q0$ estimates higher than the real fundamental scattering vector, resulting in lower estimates of D-period when not accounting for this factor (Extended Data Fig. 6b).

Statistically, D-periods estimated using 2D SAXS are reduced upon compressive loading (Extended Data Fig. 6ai) by between 0.08 – 0.276 nm (Kruskal-Wallis $H$ = 3.857, $p$ = 0.04). This is not seen in *TomoSAXS* D-period estimates, that show apparently non-significant changes upon loading (Kruskal-Wallis $H = 0.048$, $p = 0.827$) despite significant strains reported per-fibre (Fig. 3fii,iii). This difference is explained by considering the relationship between initial per-fibre D-periods to their D-period strain (Fig. 3fiii), which is roughly equal around 0 % strain and suggests that nanoscale strain in individual fibres is related to their pre-strain. A consistent tendency for fibres under tensional pre-strain to compress under load and vice versa may result in a masking of gross trends seen in 2D SAXS data when measuring properties of individual fibres.

Differences in variation in D-period are less pronounced than mean D-period when comparing 2D SAXS to *TomoSAXS* measurements. While the ability of *TomoSAXS* to deconvolute scatter across individual frames provides higher proportions of low variation estimates (Extended Data Fig. 6aiv, bi), standard deviations of measured D-period variation are not significantly different between the techniques (Kruskal-Wallis $H = 3.692$, $p = 0.051$) (Extended Data Fig. 6aii,aiv). Estimated D-period variation increase upon loading when measured using 2D SAXS (Extended Data Fig. 6aii) (Kruskal-Wallis $H = 4.857$, $p = 0.03$). As for D-period, D-period variation also shows no clear trend upon load when estimated using *TomoSAXS*. This can also be explained when considering the relationship between initial per-fibre D-period variation and change in variation upon loading (Extended Data Fig. 4d), which may also mask overarching trends.

We further characterise these differences observed in analysis of real data when measured using 2D SAXS and *TomoSAXS*, by applying these techniques for digital phantom voxels created using 3D diffraction modelling comprising three fibres of known, randomised orientation and scattering parameters (Extended Data Fig. 6b-c). Comparison of resultant estimates show that both methods produce estimates that correlate with known D-period (Extended Data Fig. 6ci) and D-period variation (Extended Data Fig. 6cii). However, *TomoSAXS* provides significantly more accurate estimates (Absolute Error; AER) for per-fibre D-period than 2D SAXS (Kruskal-Wallis $H = 29.54$, $p < 0.001$) and D-period variation (Kruskal-Wallis $H = 45.44$, $p < 0.001$). Precision (measured as standard deviation of error; SDER) is also higher for *TomoSAXS* estimations of D-period (*TomoSAXS* SDER = 1.073 nm; 2D SAXS SDER = 2.091 nm) and D-period variation (*TomoSAXS* SDER = 0.523 nm; 2D SAXS SDER = 1.061 nm). *TomoSAXS* estimates also conform closer to a slope of 1:1 with known values. The Regression between known D-period values and those estimated by 2D SAXS suggest that this method over estimates low D-period fibres and underestimates high D-period fibres (Extended Data Fig. 6ci). For D-period variation, 2D SAXS consistently overestimates known D-period variation (Extended Data Fig. 6cii).

**Supplementary Note 20 – Comparison between SAXS tensor tomography and *TomoSAXS* analysis of IVD**

*TomoSAXS* is a new contribution to a significant suite of existing techniques for tomographic analysis of SAXS data. One of the most prominent of these is SAXS Tensor Tomography (SAXS-TT). SAXS-TT was originally designed to estimate the orientation of fibres within single voxels[6]. More recent studies have applied its core spherical harmonic functions across series of sequential “shells” across $q$ to estimate and analyse intensity patterns across $q$ (Extended Data Fig. 8 ai,ii). For each shell, intensity distributions across $\chi$ are modelled, and model amplitudes used to integrate intensity across $q$ (Extended Data Fig. 8aiii). D-period and variation in D-period can be estimated from these integrations by fitting with Gaussian models to estimate $q_0$ (Equation 6) $w_a$ (Equations 7-9).

We have here attempted to compare the performance of SAXS-TT and *TomoSAXS* for estimating per-fibre D-period and D-period variation, through application of these techniques upon digital phantoms created using 3D diffraction modelling comprising of three fibres with known, randomised orientations and scattering parameters. SAXS-TT was simulated by simulating scattering intensity across $\chi$ for 200 sequential $q$ shells between 0.26-0.35 $nm^{-1}$. Each shell was fitted with a spherical harmonic function, with the estimated amplitude of the function saved to provide the simulated intensity value across the harmonic for the respective $q$ value. These values were fitted across $q$ using a Gaussian model, to provide estimates of average D-period and D-period variation across the voxel. Fibres in each voxel were also fitted using *TomoSAXS* (Extended Data Fig. 8aiv).

Comparison of estimates provided by SAXS-TT and *TomoSAXS* for D-period (Extended Data Fig. 6bi) and D-period variation (Extended Data Fig. 6bii) suggest that *TomoSAXS* provides significantly more accurate estimates than SAXS-TT. Comparisons of AER are significantly less for *TomoSAXS* than SAXS-TT for both D-period (Extended Data Fig. 8bi) (Kruskal-Wallis $H = 49.15$, $p < 0.001$) and D-period variation (Extended Data Fig. 8bii) (Kruskal-Wallis $H = 144.3$, $p < 0.001$). Precision of estimates are also greater for *TomoSAXS* compared to SAXS-TT for D-period (*TomoSAXS* SDER = 0.769 nm; SAXS-TT SDER = 1.796 nm) and D-period variation (*TomoSAXS* SDER = 0.362 nm; SAXS-TT SDER = 2.140 nm).

**Supplementary Note 21 – HiP-CT of human IVD**

Whole-segment imaging was performed at the BM18 beamline of the European Synchrotron Radiation Facility (ESRF) using hierarchical phase-contrast tomography[7] with a polychromatic beam. A combination of sapphire, silver, and glassy-carbon attenuators was used to tailor the spectrum to an average energy of 99.8 keV. The distance between the X-ray source and the sample was 178 m, while the propagation distance was 10 m. Data was acquired in continuous-rotation helical mode over 360° with half-acquisition geometry to extend the lateral field of view to 81.29 mm. A total of 15,000 projections were recorded per full rotation using an IRIS-15 (Teledyne Photometrics, United-States) camera (5,056 x 2,960 pixels) combined with a LuAG:Ce 2000 µm scintillator (custom-made by Crytur, Czechia) at an effective isotropic voxel size of 10.045 µm. Exposure time was 13 ms per projection, with 5 mm vertical translation per rotation, resulting in an axial overlap of 50%. The total scan time for the overview was 214.7 minutes. The dose rate and the total integrated surface dose were 6.9 Gy/s and 10.4 kGy, respectively. Flat- and dark-field corrections were calculated from a dedicated reference scan acquired above the sample in the formalin-filled container, under identical beam and geometry conditions.

A high-resolution local zoom was subsequently performed on a selected region of interest encompassing the central nucleus pulposus and adjacent annulus fibrosus. The beam energy was increased to 111 keV. The propagation distance was reduced to 4 m. The isotropic voxel size was 4.253 µm and the exposure time was 28 ms exposure per projection. Three contiguous z-series scans were performed with 8 mm vertical steps, covering a total axial range of 16 mm. The total acquisition time for the zoom dataset was 462.1 minutes. The dose rate and the total integrated surface dose were 1.13 Gy/s and 1.3 kGy, respectively. Reference flat-field scans were acquired before and after the z-series using the same attenuator configuration, enabling virtual flat-field correction by angular sector. For more information on the protocol see Brunet et al[8].

All projection data were reconstructed using Nabu, deployed on the ESRF high-performance computing cluster. Single-distance Paganin phase retrieval[9] was applied combined with an unsharp mask, followed by filtered back-projection. The center of rotation was determined automatically. Post-processing included, 16-bit conversion, cropping, and conversion to JPEG2000 for storage. All scan, reconstruction, and processing parameters were recorded in standardized JSON metadata files.

**Supplementary Note 22 – TomoSAXS of human IVD**

Correlative high-energy tomographic SAXS was performed on the IVD samples at the high-energy beamline ID31 at ESRF, Grenoble. Beam energy was 75 keV, exposure time 20ms and step size for scan was 100 μm. Scan sizes were taken large enough to capture the entire IVD laterally, and both IVD and part of the end plate vertically (dimensions lateral: 62 mm × vertical: 7 mm).
